# Maintenance DNA methylation is required for induced regulatory T cell reparative function following viral pneumonia

**DOI:** 10.1101/2025.02.25.640199

**Authors:** Anthony M. Joudi, Jonathan K. Gurkan, Qianli Liu, Manuel A. Torres Acosta, Kathryn A. Helmin, Luisa Morales-Nebreda, Nurbek Mambetsariev, Carla Patricia Reyes Flores, Hiam Abdala-Valencia, Elizabeth M. Steinert, Samuel E. Weinberg, Benjamin D. Singer

## Abstract

FOXP3+ natural regulatory T cells (nTregs) promote resolution of inflammation and repair of epithelial damage following viral pneumonia-induced lung injury, thus representing a cellular therapy for patients with acute respiratory distress syndrome (ARDS). Whether in vitro induced Tregs (iTregs), which can be rapidly generated in substantial numbers from conventional T cells, also promote lung recovery is unknown. nTregs require specific DNA methylation patterns maintained by the epigenetic regulator, ubiquitin-like with PHD and RING finger domains 1 (UHRF1). Here, we tested whether iTregs promote recovery following viral pneumonia and whether iTregs require UHRF1 for their pro-recovery function. We found that adoptive transfer of iTregs to mice with influenza virus pneumonia promotes lung recovery and that loss of UHRF1-mediated maintenance DNA methylation in iTregs leads to reduced engraftment and a delayed repair response. Transcriptional and DNA methylation profiling of adoptively transferred UHRF1-deficient iTregs that had trafficked to influenza-injured lungs demonstrated transcriptional instability with gain of effector T cell lineage-defining transcription factors. Strategies to promote the stability of iTregs could be leveraged to further augment their pro-recovery function during viral pneumonia and other causes of ARDS.

## Introduction

Regulatory T cells (Tregs) are a subset of CD4^+^ T cells that prevent spontaneous autoimmunity and mitigate exuberant immune responses by promoting self-tolerance and anergy (1,2). Tregs also promote repair in multiple tissue types, including the lungs, following acute injury due to infection or sterile triggers (3–11). To maintain their identity and function, Tregs require expression of the lineage-determining transcription factor, FOXP3, and maintenance of a signature DNA methylation landscape at key genomic elements, such as the *Foxp3* super-enhancer and other Treg-specific super-enhancers (Treg-SE) (12–14). Natural regulatory T cells (nTregs) emigrate from the thymus expressing FOXP3 and with the Treg lineage-determining DNA methylation landscape in place (12,13). Unfortunately, these cells are relatively rare—5–15% of circulating CD4^+^ T cells—and difficult to expand ex vivo, posing a barrier to therapeutic Treg transfer protocols (15). Induced regulatory T cells (iTregs) derive from conventional (FOXP3^−^) CD4^+^ T cells that express FOXP3 after culture in the presence of TGF-β, IL-2, and T cell receptor (TCR) stimulation, resulting in gain of nTreg-like suppressive functions. As iTregs are derived in vitro from conventional CD4^+^ T cells, a more abundant cell type, they can expand more robustly and rapidly than nTregs, an attractive feature for clinical use. Whether iTregs function like nTregs to promote recovery following acute lung injury is not known. Moreover, iTregs do not carry the signature DNA methylation landscape seen in nTregs, leading to transcriptional instability in inflammatory microenvironments and potential conversion to pro-inflammatory T cell phenotypes (12,16–18). Indeed, modulation of the epigenetic landscape via DNA methyltransferase inhibition or TET enzyme activation augments the stability and function of natural and induced Tregs (8,19–25). Elucidating targetable mechanisms that stabilize iTreg transcriptional programs and function therefore represents an important objective for developing Treg-based therapies that promote recovery from severe pneumonia (15,26).

The epigenetic regulator, ubiquitin-like with PHD and RING finger domains 1 (UHRF1, also known as Np95 in mice and ICBP90 in humans), is a multidomain nonredundant adapter protein that recruits the maintenance DNA methyltransferase, DNMT1, to replicating daughter DNA strands to maintain cell type-specific methylation patterns during DNA replication (27–29). UHRF1 is required for the stability of nTreg identity; loss of UHRF1 in nTregs during thymic development or in the adult mouse leads to generation of inflammatory ex-FOXP3 (i.e., ex-Treg) cells (30). In contrast, the necessity of UHRF1 in regulating iTreg stability and function is less clear. Published data suggest that UHRF1 is dispensable for iTreg generation (30,31), yet others have reported that iTreg generation from UHRF1-deficient conventional CD4^+^ T cells augments their suppressive function in a colitis model of inflammation (31). We hypothesized that iTregs require UHRF1-mediated maintenance DNA methylation to stabilize their acquired transcriptional and functional programs.

To test our hypothesis, we performed adoptive transfer of UHRF1-sufficient or -deficient iTregs into Treg-depleted mice with influenza A virus pneumonia, which do not recover from lung injury in the absence of reconstitution of the Treg population. Adoptive transfer of UHRF1-sufficient iTregs promoted recovery similar to adoptive transfer of nTregs. In contrast, we found that recipients of UHRF1-deficient iTregs suffered worsened hypoxemia and mortality as well as delayed alveolar epithelial repair compared with mice that received UHRF1-sufficient iTregs. UHRF1-deficient iTregs displayed reduced lung engraftment at early and late recovery timepoints. Loss of UHRF1-mediated maintenance DNA methylation had no effect on FOXP3 induction or stability yet caused significant transcriptomic instability at other core Treg loci in vitro and in vivo during viral pneumonia. In summary, UHRF1-mediated maintenance DNA methylation stabilizes iTreg cellular identity and reparative function following viral pneumonia.

## Results

### nTregs and iTregs promote recovery following viral pneumonia

Transient depletion of FOXP3^+^ nTregs is followed by a renewal of the FOXP3^+^ population (32). To determine the time window following viral infection during which FOXP3^+^ Tregs promote recovery and optimize the timing of Treg adoptive transfer, we assessed whether the timing of Treg renewal determines recovery phenotypes following influenza A virus pneumonia. To deplete Tregs, we administered loading doses of diphtheria toxin (DTx) to *Foxp3^GFP-DTR^*mice two days prior to sublethal influenza A virus infection and continued DTx administration every two days until pre-specified days post infection (DPI: 6, 10, 14, and 21) **(Supplemental Figure 1A)**. We previously reported that DTx administration to wild-type mice with viral pneumonia does not contribute to immunopathology (33). Here, we confirmed depletion of Tregs in spleen and lung tissue at 6 and 13 DPI in *Foxp3^GFP-DTR^*mice **(Supplemental Figures 1B and 1C)**. We found that *Foxp3^GFP-DTR^*mice that had DTx withdrawn at 6 DPI recovered their mass faster than mice that continued to receive DTx through days 10, 14, or 21 **(Supplemental Figure 1D)**. As administration of DTx results in some degree of lost mass (34), we examined lung injury at 60 DPI. While all groups had evidence of residual lung injury, it was most severe in mice that continued to receive DTx through 21 DPI **(Supplemental Figures 1E)**. In response to antigen or inflammation in vivo, some conventional (FOXP3^−^) CD4^+^ T cells transiently express FOXP3 but not the signature DNA methylation pattern characteristic of nTregs; these cells are known as peripheral Tregs (pTregs) (35–39). Accordingly, we determined the DNA methylation profile of FOXP3^+^ cells that renew following withdrawal of DTx in the influenza virus pneumonia model. Genome-wide DNA methylation profiling revealed that the Treg-SE DNA methylation profile of the renewed FOXP3^+^ population exhibited an nTreg-type profile **(Supplemental Figure 1F)**.

**Figure 1:**
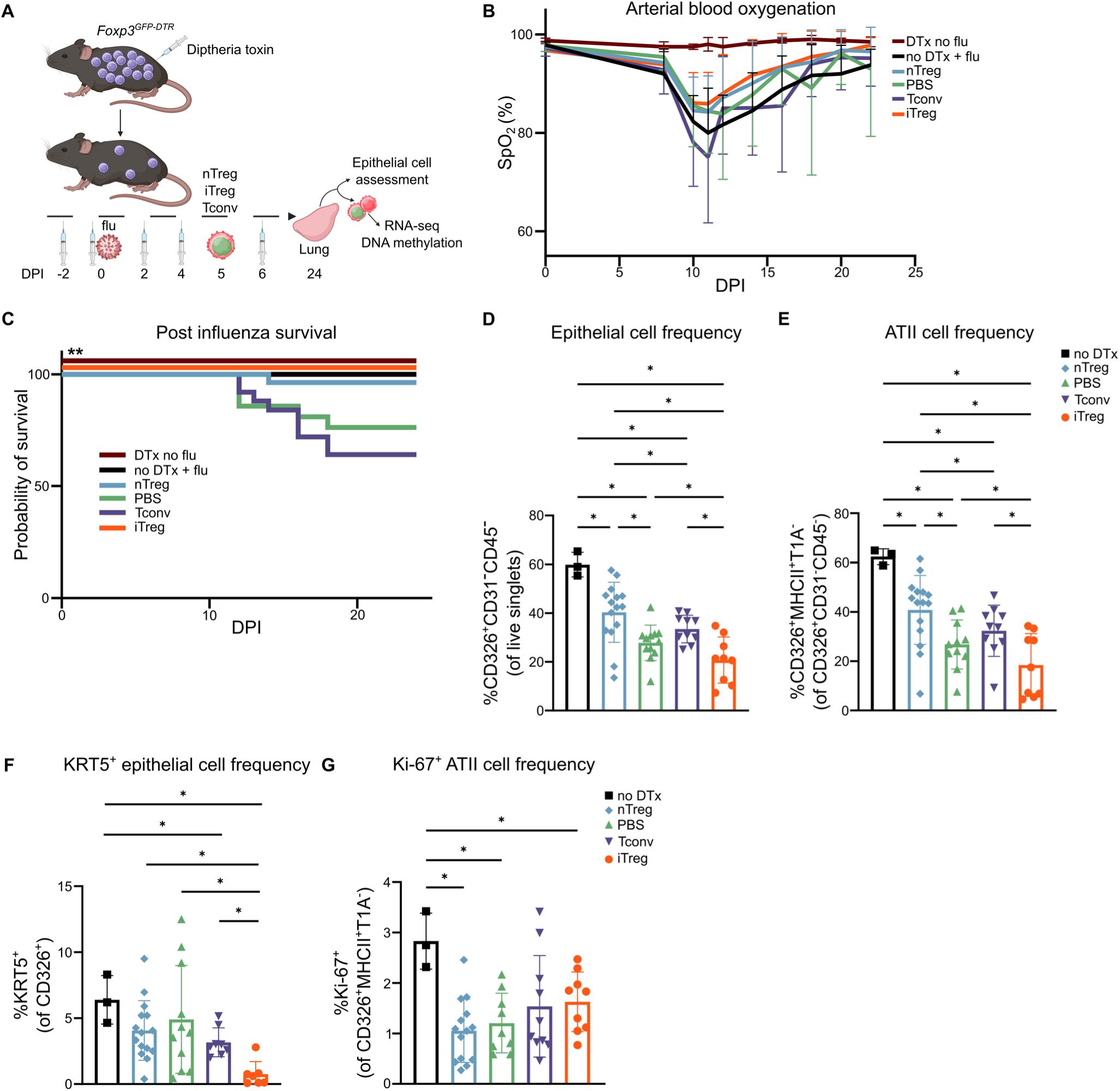
Adoptive transfer of iTregs promotes survival following viral pneumonia. **(A)** Schematic of experimental design. **(B)** Arterial oxyhemoglobin saturation (SpO_2_) over time in *Foxp3^GFP-DTR^* mice that received adoptive transfer of nTreg (n = 11), PBS (n = 9), Tconv (n = 12), or iTreg (n = 18) 5 days following intra-tracheal inoculation of 6.5 PFUs of influenza A/WSN/33 H1N1 virus. Negative controls included mice that received diphtheria toxin but no influenza (DTx no flu, n = 4) and influenza but no diphtheria toxin (no DTx + flu, n = 6). **(C)** Survival of *Foxp3^GFP-DTR^* mice that received adoptive transfer of nTreg (n = 27), PBS (n = 21), Tconv (n = 25), or iTreg (n = 18) at 5 DPI. Negative controls included no DTx + flu (n = 9) and DTx no flu (n = 4). **(D-G)** Frequency of CD326^+^CD31^−^CD45^−^ cells (D), CD326^+^MHCII^+^T1A^−^ cells (E), KRT5^+^CD326^+^ cells (F), and Ki-67^+^CD326^+^MHCII^+^T1A^−^ cells (G), at day 24 DPI in mice that received no DTx (n = 3), nTreg (n = 15), PBS (n = 11), Tconv (n = 10), or iTreg (n = 9). Survival curve (C) *p*-value was determined using the log-rank (Mantel-Cox) test, \*\**p* < 0.005. Data presented as mean and SD with * *q* < 0.05 according to multiple Mann-Whitney tests and correcting for multiple comparisons using the two-stage linear step-up procedure of Benjamini, Krieger, and Yekutieli with Q = 5% (D-G).

**Supplemental Figure 1:**
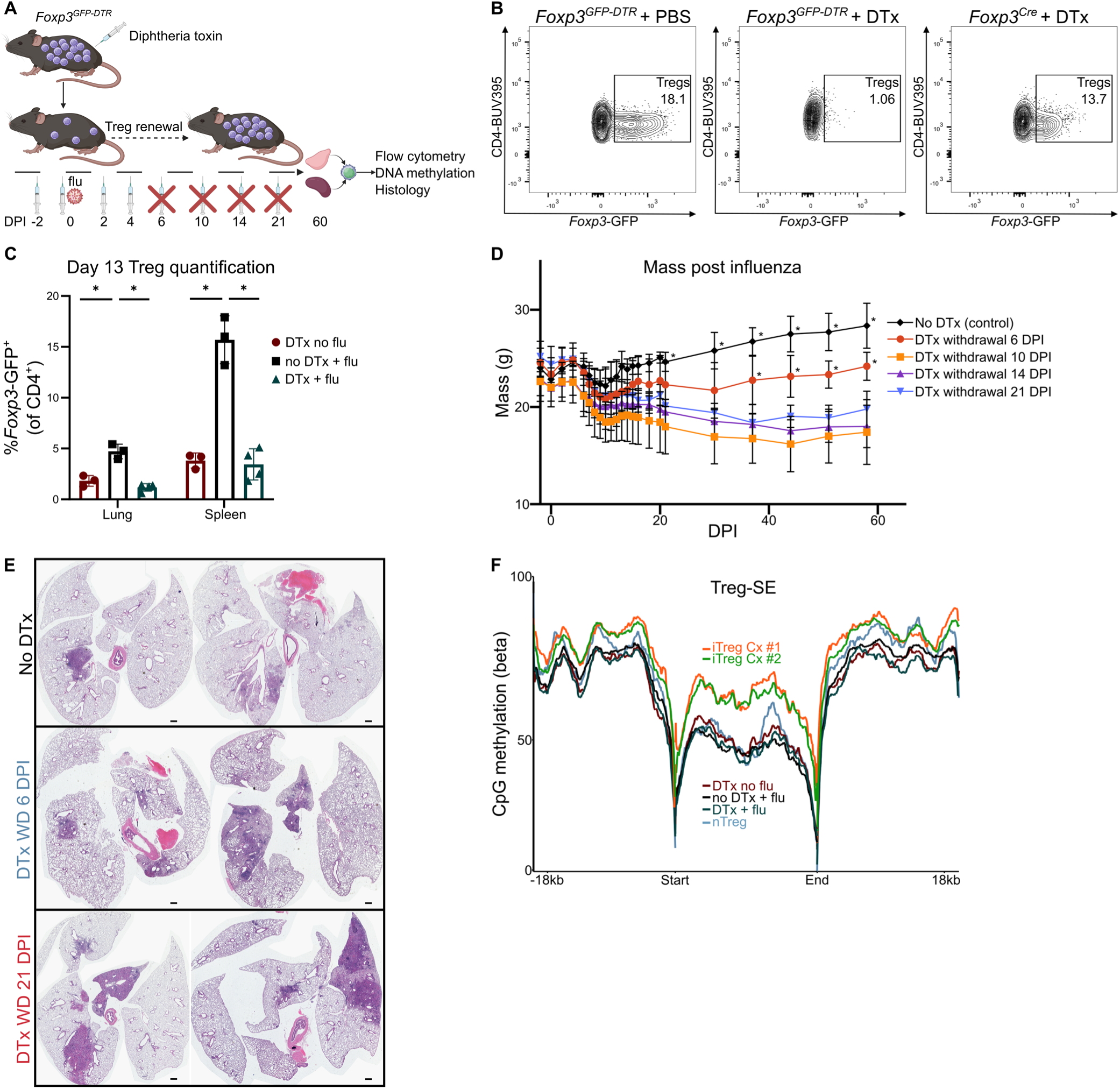
Effect of Treg renewal on recovery following influenza A virus infection. **(A)** Schematic of experimental design. **(B)** Flow cytometry contour plots demonstrating the loss of the endogenous splenic CD4^+^*Foxp3*-GFP^+^ Treg population in *Foxp3^GFP-DTR^* mice following administration of diphtheria toxin (DTx). **(C)** Lung and spleen *Foxp3*-GFP^+^ Treg cell quantification in *Foxp3^GFP-DTR^* mice that received influenza A virus but no DTx (No DTx + flu, n = 3) or influenza and DTx (DTx + flu, n = 4) at 13 DPI compared with mice that received DTx but no influenza (DTx No flu, n = 3). **(D)** Mass over time of *Foxp3^GFP-DTR^* mice receiving no DTx or DTx until pre-specified timepoints (DTx withdrawal) following intra-tracheal inoculation of 5 plaque-forming units (PFU) of influenza A/WSN/33 H1N1 virus (n = 5 per group). **(E)** Representative lung histopathology (H&E staining) at 60 DPI of control (No DTx) and DTx withdrawal mice following influenza A virus infection. Original magnification x10, scale bar = 1 mm. **(F)** Metagene analysis of DNA methylation across Treg-SE of Tregs recovered from the mice described in 1D compared with splenic nTregs and iTregs from culture. Data presented as mean and SD. * *q* < 0.05 according to one-way ANOVA with two-stage linear step-up procedure of Benjamini, Krieger, and Yekutieli with Q = 5% (C).* *q* < 0.05 according to mixed-effects model (REML) with two-stage linear step-up procedure of Benjamini, Krieger, and Yekutieli with Q = 5% (D).

We generated iTregs from mice harboring a tamoxifen-inducible *Foxp3*-Cre driver with a green fluorescent protein (GFP) label (*Foxp3^GFP-Cre-ERT2^*) and a *loxP*-flanked stop codon upstream of the red florescent protein, tdTomato, driven by a CAG promoter at the open *ROSA26* locus (*ROSA26Sor^CAG-tdTomato^*) (30). We flow cytometry sorted splenic and lymph node CD4^+^ conventional T cells (Tconv, CD4^+^*Foxp3*-GFP^−^) from *Foxp3^GFP-CreERT2^Rosa26Sor^CAG-tdTomato^* mice and cultured them in the presence of T cell receptor stimulation (αCD3/αCD28), TGF-β, IL-2, and tamoxifen to induce FOXP3^+^ cell-specific GFP and tdTomato expression **(Supplemental Figure 2A)**. Concurrently, *Foxp3^GFP-DTR^* mice received DTx followed by intratracheal instillation with a titer of influenza A virus sufficient to cause 10–20% mortality in Treg-depleted animals. At 5 DPI, 1×10^6^ iTregs, nTregs, or Tconv cells or PBS were administered via retroorbital injection **(Figure 1A)**. Although arterial oxyhemoglobin saturation (SpO_2_) was similar between groups, mice that received influenza but no DTx, nTregs (positive control), or iTregs experienced significantly greater survival compared with mice that received Tconv cells (negative control) or PBS (vehicle control) **(Figures 1B and 1C)**. Mice that received influenza but no DTx displayed a more rapid recovery in mass as well as a significantly lower absolute number of lung-infiltrating leukocytes compared with the other groups **(Supplemental Figures 2B and 2C)**. Flow cytometry analysis of lung single-cell suspensions at 24 DPI revealed a greater percentage of alveolar epithelial cells, including alveolar epithelial type 2 (ATII) cells (CD326^+^MHCII^+^T1A^−^/CD326^+^CD31^−^CD45^−^) (40), in mice that received nTregs or no DTx compared with the Tconv and PBS groups **(Figure 1D and 1E)**. Unexpectedly, mice that received iTregs displayed the lowest percentage of ATII cells among the experimental groups. To further characterize the effect of iTreg adoptive transfer on epithelial repair, we examined the KRT5+ epithelial cell population, a marker of dysregulated and incomplete repair (41–43). Mice that received iTregs displayed the lowest percentage of KRT5^+^ epithelial cells, suggesting effective repair despite the reduced percentage of ATII cells **(Figure 1F)**. We observed no significant differences in the frequency of proliferating (Ki-67^+^) ATII cells between groups that received DTx **(Figure 1G)** or in absolute epithelial or endothelial cell numbers between groups **(Supplemental Figures 2D-2H)**. Taken together, these results suggest a beneficial effect of iTregs in promoting alveolar epithelial repair.

**Figure 2:**
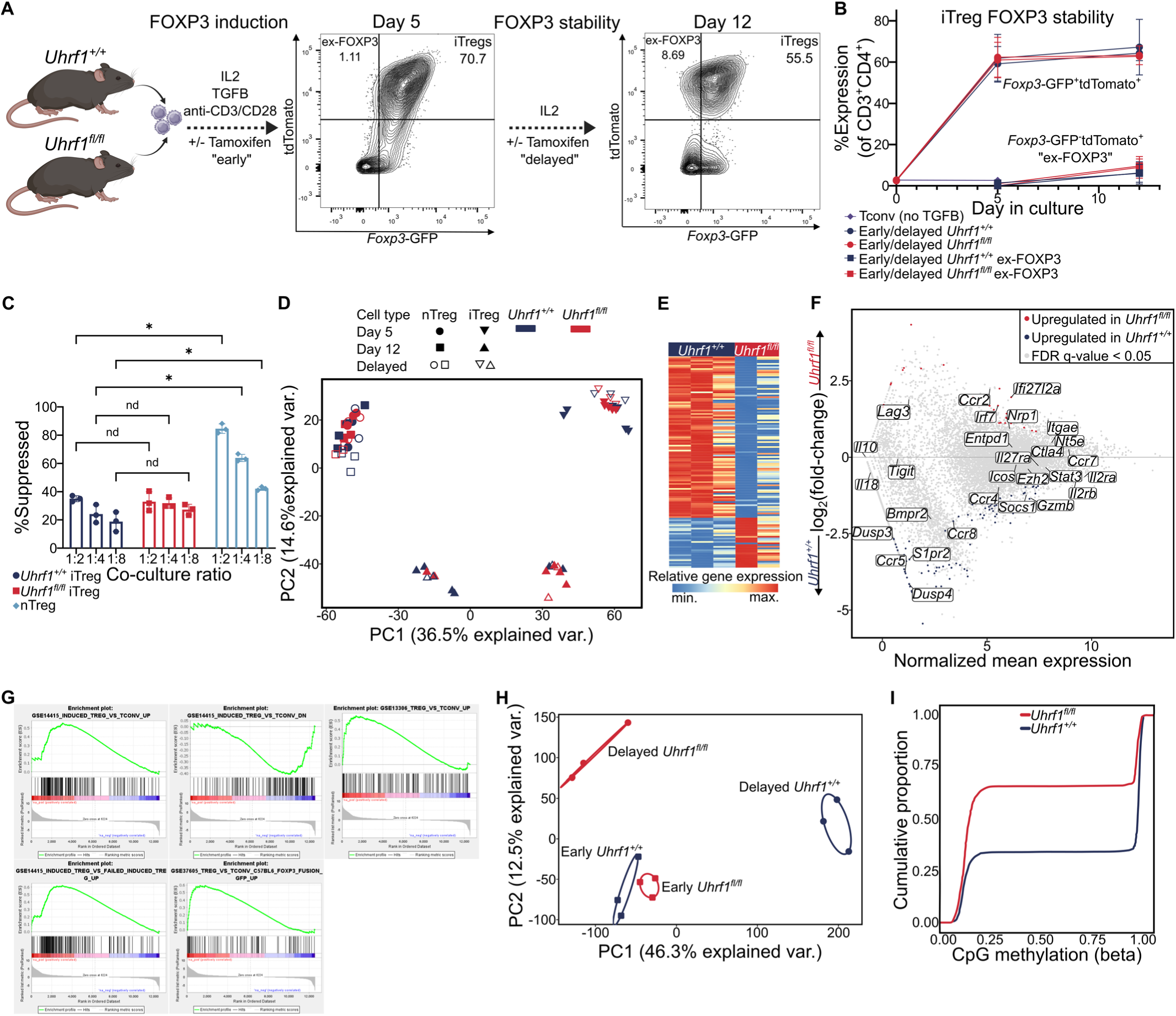
UHRF1 is dispensable for iTreg FOXP3 expression and suppressive capacity but is required for transcriptional and epigenetic stability in vitro. **(A)** Schematic of experimental design. **(B)** Frequency of *Foxp3*-GFP^+^ and *Foxp3*-GFP^−^tdTomato^+^ (ex-FOXP3) cells in culture of iTregs with UHRF1 deleted concurrent with (early) or 5 days after (delayed) FOXP3 induction compared with *Uhrf1^+/+^* controls on days 5 and 12 of culture (n = 3 per group). **(C)** Division index of CD4^+^CTV^+^*Foxp3*-GFP^−^ splenic responder T cells co-cultured for 72 hours with varying ratios of *Uhrf1^+/+^*CD4^+^*Foxp3*-GFP^+^ iTregs, *Uhrf1^fl/fl^*CD4^+^*Foxp3*-GFP^+^ iTregs, or *Uhrf1^+/+^* nTregs (n = 3 per group). **(D)** Principal component analysis of 6,978 differentially expressed genes from sorted cells described in A, identified from ANOVA-like testing with FDR *q* < 0.05. Ellipses represent normal contour lines with 1 standard deviation probability. **(E)** K-means clustering of 127 genes with an FDR *q* < 0.05 comparing the *Uhrf1^fl/fl^* and *Uhrf1^+/+^* iTreg populations on day 12 in which UHRF1 was deleted after FOXP3 induction (delayed) with k = 2. **(F)** MA plot comparing gene expression of delayed *Uhrf1^+/+^* and *Uhrf1^fl/fl^*iTregs on day 12 of culture. Genes of interest are annotated. **(G)** Enrichment plot of the GSE14415_INDUCED_TREG_VS_TCONV_UP, GSE14415_INDUCED_TREG_VS_TCONV_DN, GSE14415_INDUCED_TREG_VS_FAILED_INDUCED_TREG_UP, GSE37605_TREG_VS_TCONV_C57BL6_FOXP3_FUSION_GFP_UP, GSE13306_TREG_VS_TCONV_UP gene sets (*p* < 0.05, FDR *q* < 0.25) generated through GSEA preranked testing of the expressed genes of delayed *Uhrf1^+/+^* iTregs (control) and delayed *Uhrf1^fl/fl^* iTregs on day 12 of culture. **(H)** PCA of 81,179 differentially methylated cytosines identified from ANOVA-like testing with FDR *q* < 0.05. Ellipses represent normal contour lines with 1 standard deviation probability. **(I)** Cumulative distribution function plot of differentially methylated cytosines expressed as β scores, with 0 representing unmethylated and 1 representing fully methylated; a shift in the cumulative distribution function up and to the left represents relative hypomethylation. \**q* < 0.05 according to two-way ANOVA with two-stage linear step-up procedure of Benjamini, Krieger, and Yekutieli with Q = 5.

**Supplemental Figure 2:**
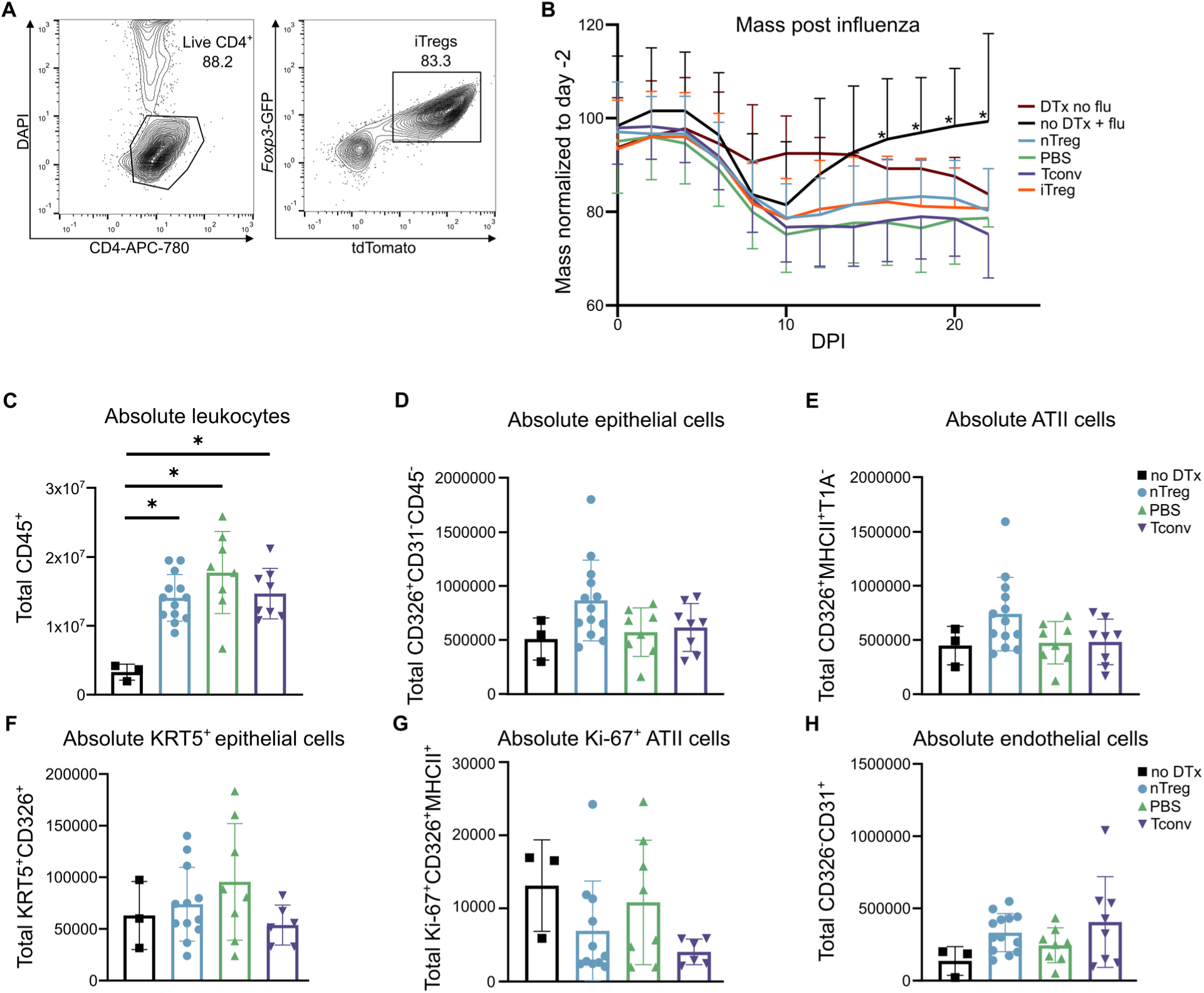
Effects of receiving DTx or no DTx. **(A)** Flow cytometry contour plots for phenotyping of the CD4^+^*Foxp3*-GFP^+^tdTomato^+^ iTreg population following culture in the presence of αCD3/αCD28, IL-2, TGF-β, and tamoxifen. **(B)** Mass over time of *Foxp3^GFP-DTR^* mice receiving adoptive transfer of nTreg (n = 27), PBS (n = 21), Tconv (n = 25), or iTreg (n = 18) cells 5 days following intra-tracheal inoculation of 6.5 PFU of influenza A/WSN/33 H1N1 virus. Negative controls included mice that received influenza but no DTx (no DTx + flu, n = 9) and DTx but no influenza (DTx no flu, n = 3). **(C)** Total number of lung CD45^+^ cells at 24 DPI in mice that received No DTx (n = 3), nTreg (n = 15), PBS (n = 11), or Tconv (n = 10). **(D-H)** Total number of lung CD326^+^CD31^−^CD45^−^ cells (D), CD326^+^MHCII^+^T1A^−^ cells (E), KRT5^+^CD326^+^ cells (F), Ki-67^+^CD326^+^MHCII^+^ cells (G), and CD326^−^CD31^+^ cells (H) at 24 DPI in mice that received No DTx (n = 3), nTreg (n = 15), PBS (n = 11), or Tconv (n = 10) cells. * *q* < 0.05 according to mixed-effects model (REML) with two-stage linear step-up procedure of Benjamini, Krieger, and Yekutieli with Q = 5% (B). Data presented as mean and SD (C, F-H) with * *q* < 0.05 according to multiple Mann-Whitney tests and correcting for multiple comparisons using the two-stage linear step-up procedure of Benjamini, Krieger, and Yekutieli with Q = 5% (C). *UHRF1 is dispensable for iTreg FOXP3 induction and stability but is required to maintain transcriptional and epigenetic programs in vitro*.

We tested whether UHRF1-mediated maintenance DNA methylation is necessary for iTreg differentiation and stability in vitro. We bred *Uhrf1^fl/fl^Foxp3^GFP-CreERT2^Rosa26Sor^CAG-tdTomato^* mice (referred to here as *Uhrf1^fl/fl^*), allowing us to generate inducible, Treg-specific, *Foxp3* lineage-traceable iTregs that lose UHRF1 expression contemporaneously with FOXP3 induction (30). CD4^+^*Foxp3*-GFP^−^ T cells were isolated from the spleens of *Uhrf1^+/+^* (control) or *Uhrf1^fl/fl^* mice and cultured in iTreg-skewing media plus tamoxifen and subsequently flow cytometry sorted for bulk RNA-seq analysis (denoted as the “early” group) **(Figure 2A)**. A separate set of cells that had not been exposed to tamoxifen during the FOXP3 induction phase were transitioned into resting media, defined by the absence of TGF-β and T cell receptor stimulation, after 5 days in culture. Tamoxifen was added to the resting media and the cells were cultured for an additional 7 days (denoted as the “delayed” group) followed by flow cytometry sorting for RNA-seq analysis after 12 total days in culture. Consistent with published data (30,31), flow cytometry analysis demonstrated no significant difference in FOXP3 induction or stability in iTregs over the culture period regardless of the timing of UHRF1 deletion **(Figure 2B)**. We confirmed that CD4^+^*Foxp3*-GFP^+^ iTregs and nTregs from *Uhrf1^+/+^* mice expressed high levels of canonical Treg cell signature genes (e.g., *Il2ra*, *Il2rb*, *Icos*, *Tigit*, *Il10*, *Gzmb*, *Ctla4*, *Nt5e*, *Itgae*, *Nrp1*, and *Lag3*) **(Supplemental Figure 3A)**. We additionally confirmed the in vitro suppressive function of iTregs following 5 days of culture, finding no significant difference between *Uhrf1^+/+^* and *Uhrf1^fl/fl^* cells **(Figure 2C)**. Principal component analysis (PCA) of 6,978 differentially expressed genes (DEGs, FDR *q* < 0.05) in *Uhrf1^+/+^* and *Uhrf1^fl/fl^*cells demonstrated clustering by cell culture condition and cell type. PC1 reflected the transcriptional differences between nTregs and iTregs; PC2 reflected time in culture (day 5 versus day 12) **(Figure 2D)**. Notably, transcriptional differences in nTregs as a function of time in culture at which UHRF1 was deleted were minimal, as clustering remained tight regardless of time in culture and timing of UHRF1 loss. The loss of UHRF1 concurrent with FOXP3 induction (early) in iTregs had a nominal effect (≤3 DEGs) on the iTreg transcriptome when compared with *Uhrf1^+/+^*iTregs. Nevertheless, within PC2, we noted sub clustering within iTregs on day 12 of culture that reflected the time when UHRF1 was deleted after FOXP3 induction. Pairwise comparison of *Uhrf1^+/+^*and *Uhrf1^fl/fl^* cells in the day 12 delayed group revealed 127 DEGs, with genes upregulated in *Uhrf1^+/+^* iTregs including those associated with chemotaxis and migration (*Ccr5*, *Ccr8*, and *S1pr2*) as well as Treg proliferation, differentiation, and transcriptional stabilization (*Skp2*, *Lif*, and *Dusp4*) **(Figure 2E-F and Supplemental Table 1)**. Gene set enrichment analysis (GSEA) comparing *Uhrf1^+/+^* with *Uhrf1^fl/fl^* iTregs revealed positive enrichment of genes previously annotated to be upregulated in nTregs and iTregs compared with CD4^+^ Tconv cells and negative enrichment in genes previously annotated to be downregulated in iTregs compared with CD4^+^ Tconv cells (44–46). This analysis also revealed positive enrichment of genes previously annotated to be upregulated following successful induction of FOXP3 compared with CD4+ Tconv cells from the same culture that failed to express FOXP3 (44). Additional GSEA demonstrated positive enrichment of hallmark processes in *Uhrf1^+/+^* cells associated with Treg function, including Myc targets, E2F targets, TGF-β signaling, WntB catenin signaling, TNF-α signaling via NFκB, MTORC signaling, KRAS signaling, and IL-2–STAT5 signaling **(Figure 2G, Supplemental Figure 3B, and Supplemental Table 2)**. No hallmark gene sets were significantly positively enriched in *Uhrf1^fl/fl^* iTregs. To confirm that maintenance DNA methylation was lost upon UHRF1 deletion, we performed genome-wide 5′-cytosine–phosphate–guanine-3′ (CpG) methylation profiling with modified reduced representation bisulfite sequencing at day 12 of culture. PCA of approximately 80,000 differentially methylated cytosines (FDR *q* < 0.05) revealed distinct clustering according to culture condition **(Figure 2H)**. PC1 reflected methylation changes based on the deletion of UHRF1, whereas PC2 reflected methylation changes based on the timing of UHRF1 deletion. Pairwise comparison of *Uhrf1^+/+^* and *Uhrf1^fl/fl^*iTregs in the delayed group at day 12 demonstrated hypomethylation in *Uhrf1^fl/fl^*iTregs. **(Figure 2I)**. Taken together, these results indicate that UHRF1-mediated maintenance DNA methylation is dispensable for the establishment of iTreg FOXP3 expression, transcriptional identity, and suppressive function but is necessary for the subsequent stability of the iTreg transcriptomic signature in vitro.

**Supplemental Figure 3:**
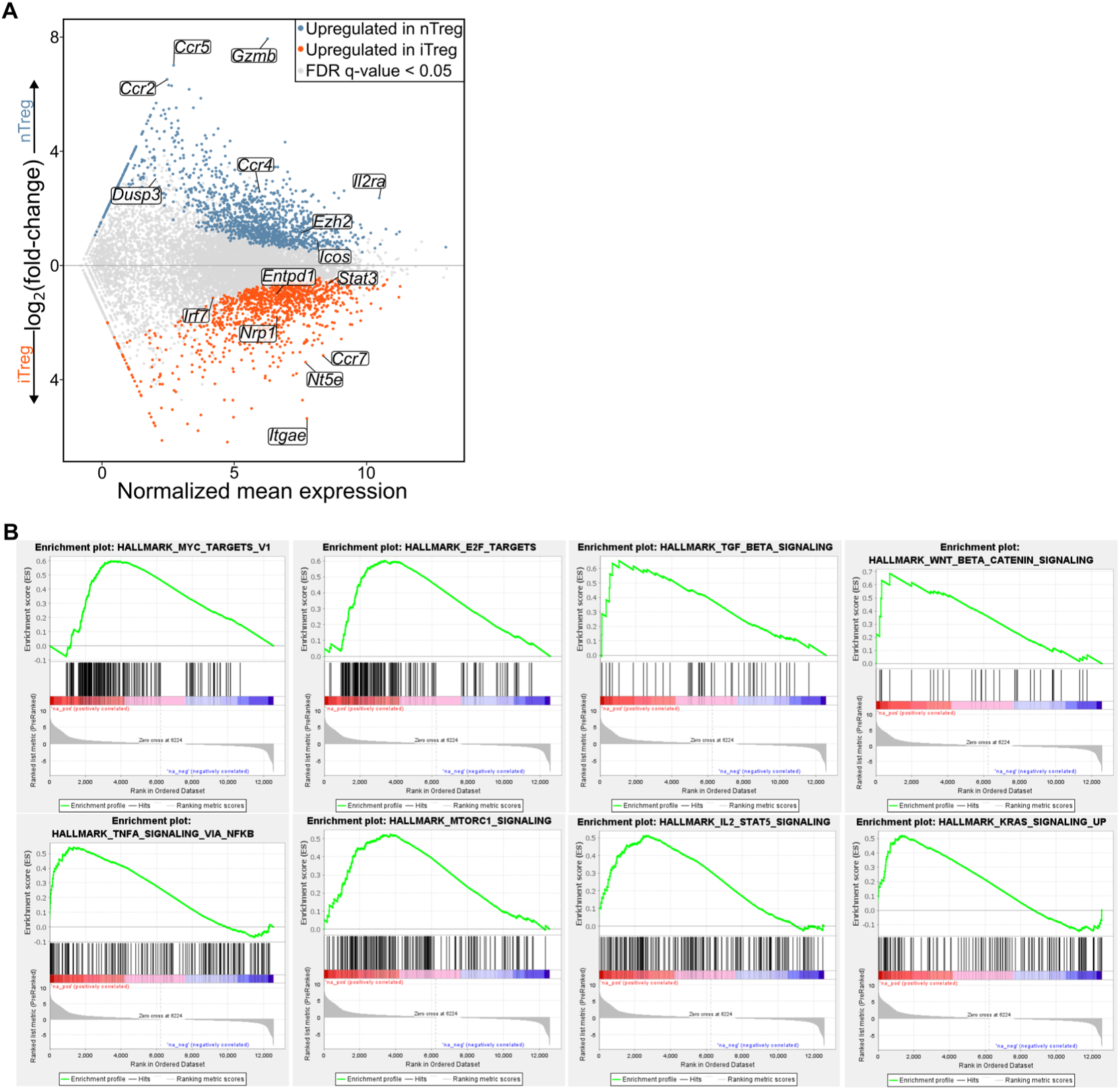
Confirmation of canonical Treg transcriptomic signature in nTregs and iTregs. **(A)** MA plot comparing gene expression of *Uhrf1^+/+^* iTregs (control) with CD4^+^*Foxp3*-GFP^+^ nTregs on day 5 of culture. Genes of interest are annotated. **(B)** Enrichment plots of the HALLMARK_MYC_TARGETS_V1, HALLMARK_E2F_TARGETS, HALLMARK_TGF_BETA_SIGNALING, HALLMARK_WNT_BETA_CATENIN_SIGNALING, HALLMARK_TNFA_SIGNALING_VIA_NFKB, HALLMARK_MTORC1_SIGNALING, HALLMARK_IL2_STAT5_SIGNALING, and HALLMARK_KRAS_SIGNALING_UP gene sets generated through GSEA preranked testing of the expressed genes of delayed *Uhrf1^+/+^* iTregs (control) and delayed *Uhrf1^fl/fl^*iTregs on day 12 of culture.

### UHRF1-deficient iTregs fail in promoting recovery following viral pneumonia

As the loss of UHRF1-mediated maintenance DNA methylation promoted transcriptional instability in iTregs, we sought to determine whether the loss of UHRF1 limits the ability of adoptively transferred iTregs to promote recovery following influenza A virus pneumonia. Using the data generated from recipients of UHRF1-sufficient iTregs *(Uhrf1^+/+^*) as a control, we found that mice that received UHRF1-deficient (*Uhrf1^fl/fl^*) iTregs experienced worsened mortality and hypoxemia compared with mice that received *Uhrf1^+/+^* iTregs **(Figure 3A-B)**. We observed no significant differences in mass recovery **(Supplemental Figure 4A).** Flow cytometry analysis of lung single-cell suspensions at 24 DPI revealed a greater frequency and total number of alveolar epithelial cells and alveolar epithelial type 2 (ATII) cells in recipients of *Uhrf1^fl/fl^* iTregs compared with recipients of *Uhrf1^+/+^*iTregs **(Figure 3C-3F)**. Notably, recipients of *Uhrf1^fl/fl^* iTregs also displayed a greater frequency and total number of KRT5^+^ epithelial cells and a higher total number of Ki-67^+^ ATII cells compared with recipients of *Uhrf1^+/+^* iTregs, suggesting a greater degree of peak injury in the recipients of *Uhrf1^fl/fl^* iTregs **(Figure 3G-3I).** Collectively, adoptive transfer of UHRF1-deficient iTregs compromised recovery from viral pneumonia and suggested a greater degree of peak injury in recipients of *Uhrf1^fl/fl^* iTregs.

**Figure 3:**
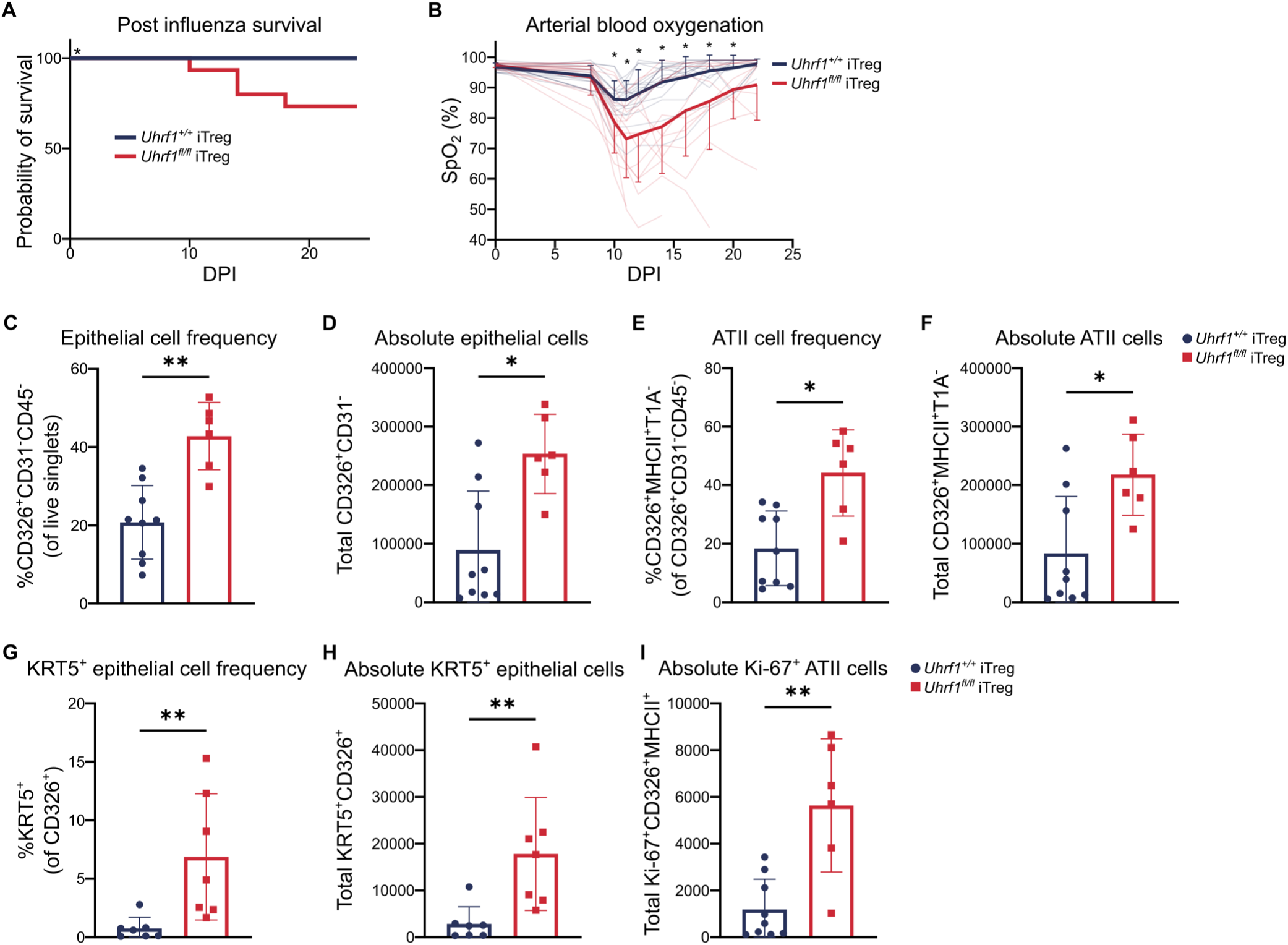
Loss of UHRF1 is sufficient to impair repair capabilities of iTregs during viral pneumonia. **(A**) Survival of *Foxp3^GFP-DTR^* mice that received *Uhrf1^+/+^* (n = 18) or *Uhrf1^fl/fl^* (n = 15) iTregs following intra-tracheal inoculation of 6.5 plaque forming units (PFUs) of influenza A/WSN/33 H1N1 virus. (**B**) Arterial oxyhemoglobin saturation (SpO_2_) over time in mice from A. **(C-F)** Frequency and total number of CD326^+^CD31^−^ cells (C-D) and CD326^+^MHCII^+^T1A^−^ cells (E-F) in the post-caval lung lobes at 24 DPI in *Foxp3^GFP-DTR^* mice that received *Uhrf1^+/+^* (n = 9) or *Uhrf1^fl/fl^* (n = 6) iTregs. **(G-I)** Frequency and total number of KRT5^+^CD326^+^ cells in the post-caval lung lobes of mice from C-F. **(I)** Total number of Ki-67^+^CD326^+^MHCII^+^ cells at 24 DPI in mice that received *Uhrf1^+/+^* (n = 9) or *Uhrf1^fl/fl^* (n = 6) iTregs. Survival curve (A) *p*-value was determined using log-rank (Mantel-Cox) test, \**p* < 0.05. \**p* < 0.05 or * *q* < 0.05 according to mixed-effects model (REML) with two-stage linear step-up procedure of Benjamini, Krieger, and Yekutieli with Q = 5% (B). Data presented as mean and SD with \**p* < 0.05 according to Mann-Whitney U test (C-I). Data from recipients of *Uhrf1^+/+^* iTregs are duplicated from results presented in Figure 1.

**Supplemental Figure 4:**
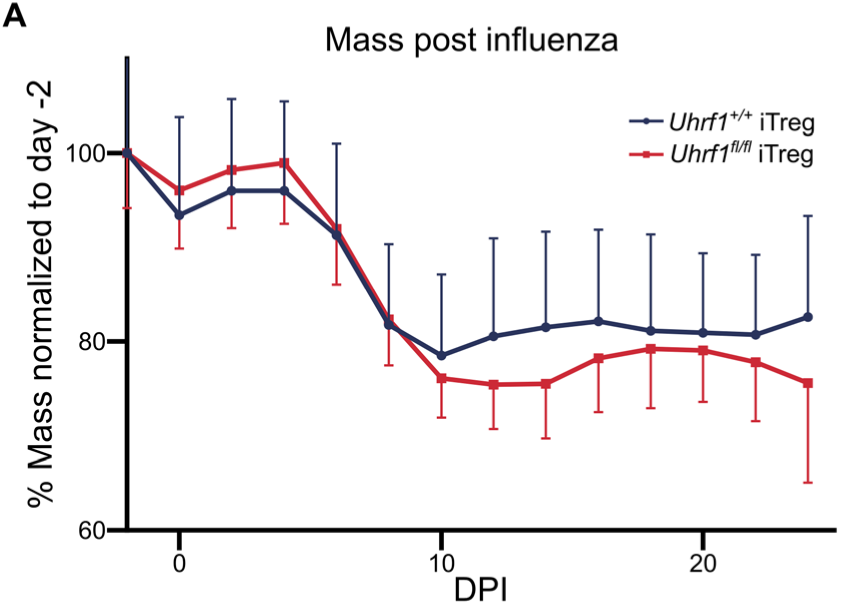
iTreg UHRF1 is dispensable for promoting mass recovery following viral pneumonia. **(A)** Mass over time of *Foxp3^GFP-DTR^* mice that received *Uhrf1^+/+^* (n = 18) or *Uhrf1^fl/fl^* (n = 15) iTregs following intra-tracheal inoculation of 6.5 PFUs of influenza A/WSN/33 H1N1 virus.

### Adoptive transfer of UHRF1-deficient iTregs results in delayed repair of lung injury following viral pneumonia compared with UHRF1-sufficient iTregs

In addition to promoting repair, Tregs possess distinct tissue-protective properties that impart resilience to damage (47). As analysis of our adoptive transfer experiments at a late repair timepoint (24 DPI) demonstrated a reduced frequency of ATII cells in recipients of *Uhrf1^+/+^* iTregs compared with recipients of *Uhrf1^fl/fl^* iTregs despite reduced hypoxemia and mortality, we hypothesized that *Uhrf1^+/+^* iTregs impart resilience to lung injury by decreasing infiltration by inflammatory immune cells or by dampening early injury to result in a less robust repair response later in the disease course. We therefore performed an additional series of adoptive transfer experiments with analysis at 11 DPI, a timepoint that correlated with peak lung injury. We found no difference in immune cell infiltration, including total leukocytes as well as myeloid and lymphoid cell subsets **(Supplemental Figure 5A-F)**. In contrast, lungs from mice that received *Uhrf1^+/+^* iTregs displayed a significantly greater total number of ATII cells with a concomitantly greater total number of Ki-67^+^ ATII cells compared with recipients of *Uhrf1^fl/fl^* iTregs **(Figure 4A-B)**. No significant differences were observed in total numbers of epithelial, KRT5^+^ epithelial, or endothelial cells (**Supplemental Figure 5G-I)**. We quantified the adoptively transferred tdTomato^+^ iTregs and found a significantly lower total number of *Uhrf1^fl/fl^* iTregs in the lungs compared with *Uhrf1^+/+^* iTregs **(Figure 4C)**. No significant difference was again observed in the frequency of ex-FOXP3 cells between groups **(Figure 4D)**. To assess whether the inflammatory microenvironment of the lung could be influencing the differences in engraftment following adoptive transfer, we quantified tdTomato^+^ iTregs from the spleens of these mice and found that the number of iTregs was lower in recipients of *Uhrf1^fl/fl^*compared with *Uhrf1^+/+^* iTregs **(Figure 4E)**. No difference was observed in the frequency of ex-FOXP3 cells **(Figure 4F)**. Taken together with the data from 24 DPI, these results suggest that UHRF1-deficient iTregs provide an insufficient early tissue-protective response that results in delayed repair.

**Figure 4:**
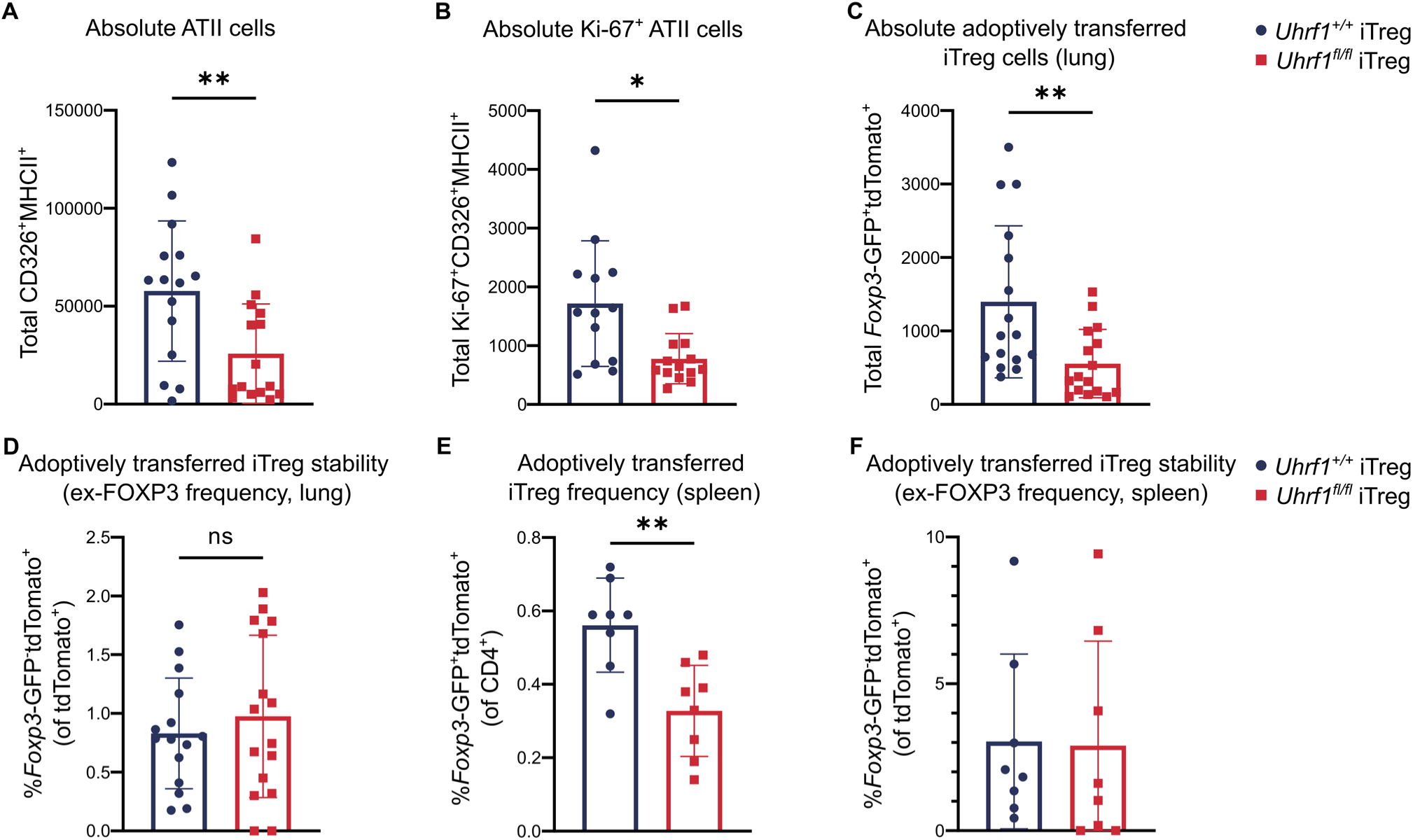
UHRF1-deficient iTregs promote an insufficient tissue-protective response during peak lung injury. **(A)** Total number of ATII cells at 11 DPI in the post-caval lobes of *Foxp3^GFP-DTR^* mice that received *Uhrf1^+/+^* iTreg (n = 15) or *Uhrf1^fl/fl^*iTreg (n = 15). **(B)** Total number of Ki-67^+^ ATII cells in the post-caval lobes of mice that received *Uhrf1^+/+^* iTreg (n = 14) or *Uhrf1^fl/fl^* iTreg (n = 14). **(C)** Total number of tdTomato^+^ cells recovered from the lungs of recipients of *Uhrf1^fl/fl^*(n = 16) or *Uhrf1^+/+^* iTregs (n = 16). **(D)** Frequency of *Foxp3*-GFP^−^tdTomato^+^ cells recovered from the lungs of recipients of *Uhrf1^+/+^* (n = 15) or *Uhrf1^fl/fl^* (n = 16) iTregs. **(E)** Frequency of tdTomato^+^ cells recovered from the spleens of recipients of *Uhrf1^+/+^* (n = 8) or *Uhrf1^fl/fl^*(n = 8) iTregs. **(F)** Frequency of *Foxp3*-GFP^−^tdTomato^+^ cells recovered from the spleens of recipients of *Uhrf1^+/+^* (n = 8) or *Uhrf1^fl/fl^* (n = 8) iTregs. Data presented as mean and SD. \**p* < 0.05; \*\**p* < 0.005; ns; not significant, according to Mann-Whitney U test **(A-E)**.

**Supplemental Figure 5:**
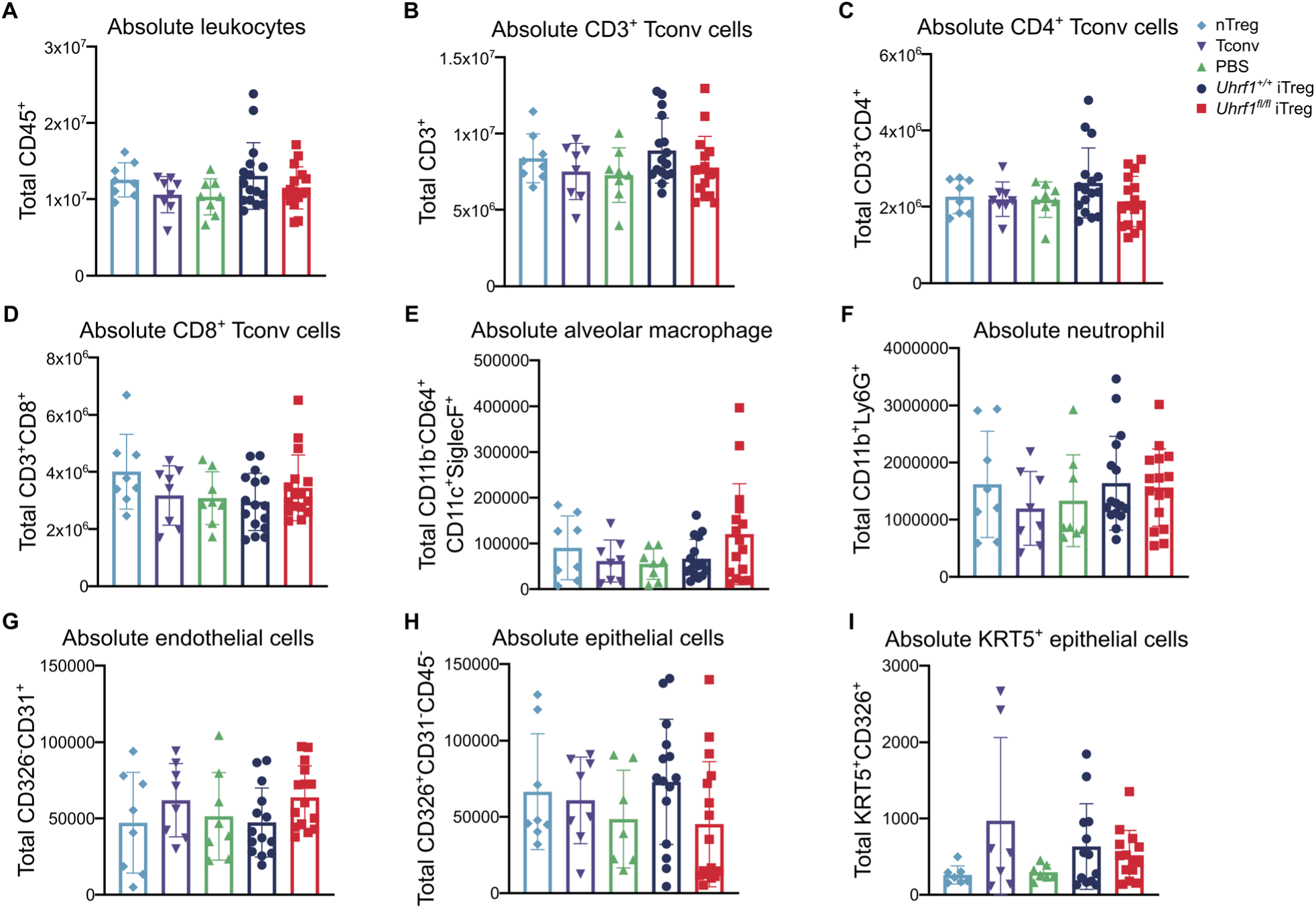
UHRF1 is dispensable for iTreg suppression of infiltrating immune cells during viral pneumonia. **(A-F)** Total numbers of CD45^+^ (A), CD3^+^ (B), CD4^+^CD8^−^ (C), CD4^−^CD8^+^ (D), CD11b^−^CD64^+^CD11c^+^SiglecF^+^ (E), and CD11b^+^Ly6G^+^ (F) cells in the lungs at 11 DPI of *Foxp3^GFP-DTR^*mice that received nTreg (n = 8), Tconv (n = 8), PBS (n = 8), *Uhrf1^+/+^* iTregs (n = 16), or U*hrf1^fl/fl^*iTregs. (n = 16) **(G)** Total number of CD326^+^CD31^−^CD45^−^ (epithelial) cells in the lungs at 11 DPI of *Foxp3^GFP-DTR^* mice that received nTreg (n = 8), Tconv (n = 8), PBS (n = 7), *Uhrf1^+/+^* iTregs (n = 15), or U*hrf1^fl/fl^* iTregs (n = 16). **(H)** Total number of KRT5^+^CD326^+^ epithelial cells in the lungs at 11 DPI of *Foxp3^GFP-DTR^* mice that received nTreg (n = 7), Tconv (n = 7), PBS (n = 7), *Uhrf1^+/+^*iTregs (n = 13), or U*hrf1^fl/fl^* iTregs (n = 14). **(I)** Total number of CD326^−^CD31^+^ (endothelial) cells in the lungs at 11 DPI of *Foxp3^GFP-DTR^* mice that received nTreg (n = 8), Tconv (n = 8), PBS (n = 8), *Uhrf1^+/+^* iTregs (n = 14), or U*hrf1^fl/fl^*iTregs (n = 15). Data presented as mean and SD. ns = not significant by multiple Mann-Whitney tests and correcting for multiple comparisons using the two-stage linear step-up procedure of Benjamini, Krieger, and Yekutieli with Q = 5% test.

### UHRF1-deficient iTregs display transcriptomic instability and poor engraftment after adoptive transfer into mice with viral pneumonia

To explore cell-intrinsic mechanisms underlying the loss of pro-recovery function in UHRF1-deficient iTregs, we profiled *Foxp3*-GFP^+^tdTomato^+^ iTregs obtained from the lungs of *Uhrf1^+/+^* and *Uhrf1^fl/fl^* iTreg recipients at 24 DPI. The frequency of adoptively transferred cells that had lost FOXP3 expression (ex-FOXP3 cells) was minimal and not significantly different between *Uhrf1^+/+^*and *Uhrf1^fl/fl^* iTregs **(Figure 5A)**. Our analysis therefore focused on transcriptomic comparison of *Uhrf1^+/+^* and *Uhrf1^fl/fl^* iTregs to identify transcriptional differences. We identified 1,187 DEGs; k-means clustering of these DEGs identified two distinct clusters **(Figure 5B and Supplemental Table 1)**. Genes upregulated in *Uhrf1^+/+^* iTregs included several associated with pro-repair function, such as

**Figure 5:**
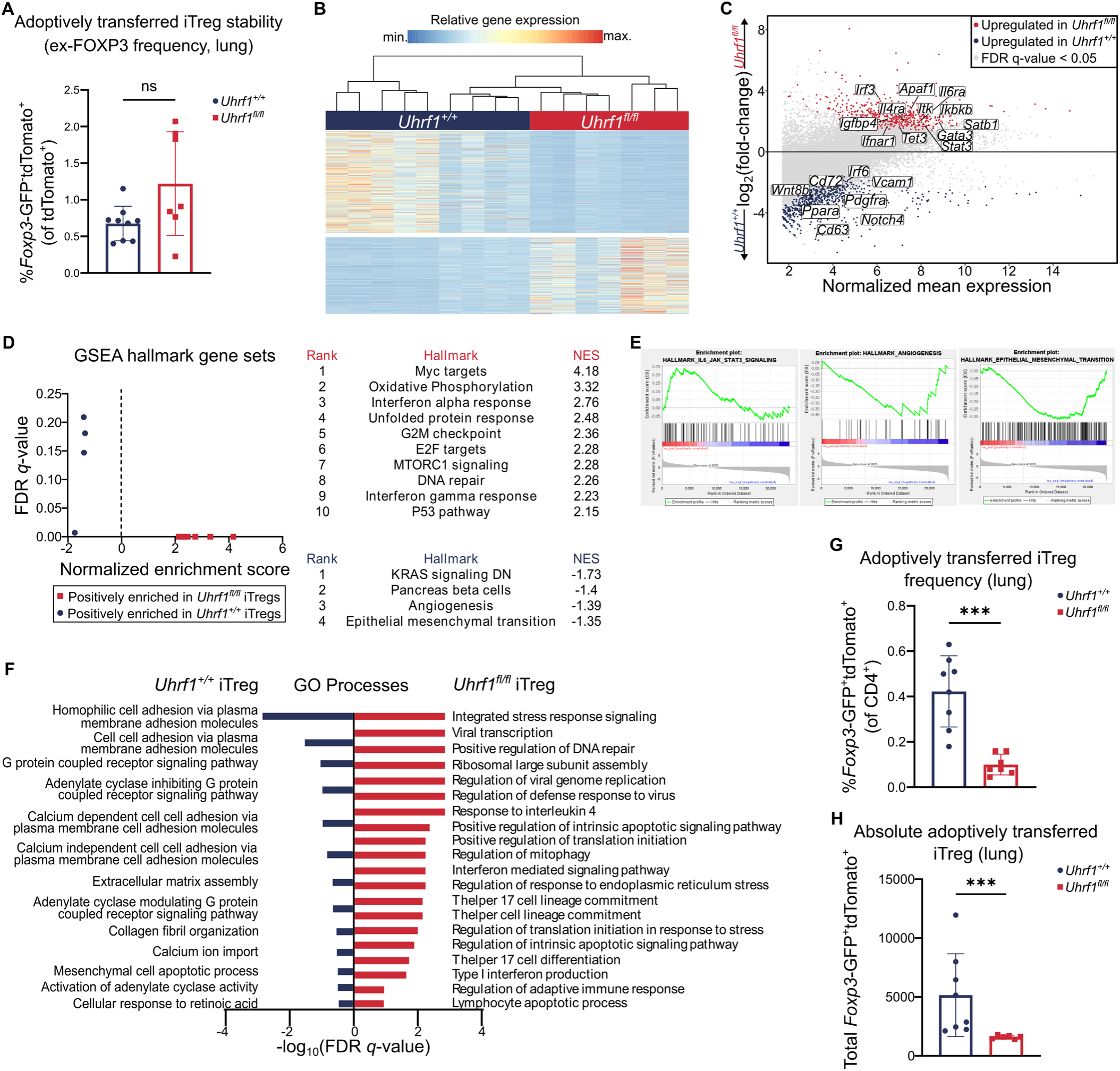
UHRF1 is required for iTreg phenotypic stability and lung tissue engraftment following viral pneumonia. **(A)** Frequency of *Foxp3*-GFP^−^tdTomato^+^ cells recovered at 24 DPI from the lungs of *Foxp3^GFP-DTR^* mice that received *Uhrf1^+/+^* (n = 9) or *Uhrf1^fl/fl^* (n = 7) iTregs. **(B)** K-means clustering of 1,187 genes with FDR *q* < 0.05 comparing adoptively transferred *Uhrf1^+/+^* and *Uhrf1^fl/fl^* iTregs at 24 DPI with k = 2. **(C)** MA plot comparing gene expression of adoptively transferred *Uhrf1^+/+^* and *Uhrf1^fl/fl^* iTregs at 24 DPI. Genes of interest are annotated. **(D)** GSEA dot plot highlighting key statistics (FDR *q*-value and normalized enrichment score or NES) and enriched gene sets. Red dots denote gene sets with a positive enrichment score or enrichment at the top of the ranked list. Blue dots denote gene sets with a negative enrichment score or enrichment at the bottom of the ranked list. **(E)** Enrichment plots of the HALLMARK_IL6_JAK_STAT3_SIGNALING, HALLMARK_ANGIOGENESIS, and HALLMARK_EPITHELIAL_MESENCHYMAL_TRANSITION gene sets generated through GSEA preranked testing of the expressed genes in *Uhrf1^+/+^* and *Uhrf1^fl/fl^* iTregs. **(F)** Selected gene ontology (GO) processes from 945 and 105 total enriched gene sets with FDR *q* < 0.25 in *Uhrf1^fl/fl^*and *Uhrf1^+/+^* iTregs, respectively. Gene sets are annotated and ranked by –log_10_-transformed FDR *q*-value. **(G)** Frequency of tdTomato^+^ iTregs recovered from the lungs at 24 DPI of mice that received *Uhrf1^+/+^* (n = 8) or *Uhrf1^fl/fl^* (n = 7) iTregs. **(H)** tdTomato^+^ iTregs recovered from the lungs at 24 DPI of mice that received *Uhrf1^+/^*^+^ (n = 8) or *Uhrf1^fl/fl^* (n = 6) iTregs. \*\*\**p* < 0.0005, ns, not significant according to Mann-Whitney U test.

*Notch4*, *Dll4*, *Pdgfra*, *Mmp12*, *Fgfr1*, *Loxl2*, *Wnt8b*, and *Yap1* **(Figure 5C)**. Notably, genes upregulated in *Uhrf1^fl/fl^*iTregs included several associated with effector helper T cell lineage commitment, such as *Stat3*, *Il6ra*, *Jak1*, *Itk*, *Igfbp4*, *Zeb1*, *Gata3*, *Il4ra*, and *Ifnar1*. GSEA revealed positive enrichment of genes associated with IL-6–STAT3 signaling as well as negative enrichment of genes associated with angiogenesis and epithelial to mesenchymal transition in *Uhrf1^fl/fl^* iTregs **(Figure 5D-E and Supplemental Table 3)**. Functional enrichment analysis revealed upregulation of gene sets associated with protein translation, interferon and interleukin signaling, helper T cell and Th17 differentiation, viral processes, DNA damage repair, cellular stress responses, and apoptotic processes in *Uhrf1^fl/fl^*iTregs **(Figures 5F and Supplemental Table 3)**. To determine the efficiency of lung engraftment of iTregs after adoptive transfer, we quantified the number of transferred iTregs in the lungs of recipient mice at 24 DPI and found a significantly reduced frequency and total number of *Uhrf1^fl/fl^* iTregs compared with *Uhrf1^+/+^* iTregs **(Figure 5G-H)**.

To assess for extra-pulmonary signatures due to loss of UHRF1, we sorted *Foxp3*-GFP^+^tdTomato^+^ and *Foxp3*-GFP^+^tdTomato^+^ (ex-FOXP3) cells from single-cell suspensions of spleens at 24 DPI for gene expression profiling. PCA of 457 differentially expressed genes identified following ANOVA-like testing with FDR *q* < 0.05 demonstrated clustering by genotype and FOXP3 expression **(Supplemental Figure 6A)**. PC1 reflected the transcriptional differences dependent on FOXP3 expression and PC2 reflected differences between genotype (*Uhrf1^+/+^*versus *Uhrf1^fl/fl^*). Pairwise comparison of *Uhrf1^+/+^*and *Uhrf1^fl/fl^* FOXP3^+^ cells revealed 183 DEGs, with genes upregulated in *Uhrf1^+/+^* iTregs associated with cell cycle regulation/cellular proliferation and induction and maintenance of Treg function (*E2f3*, *Ncoa3*, *Hpse*) **(Supplemental Figure 6B-C and Supplemental Table 1).** Genes upregulated in *Uhrf1^fl/fl^* iTregs included those associated with maintenance of function but also proinflammatory cytokines and cytokine receptors (*Mst1*, *Tmed4, Il1b*, *Il17rb*, and *Il4*). Pairwise comparison of *Uhrf1^+/+^* and *Uhrf1^fl/fl^*ex-FOXP3 cells revealed 274 DEGs, with genes upregulated in *Uhrf1^+/+^*ex-FOPX3 cells associated with Treg stability, suppressive function (*Ikzf2, Zap70, Tnfrsf9, Il1r1,* and *Parp11*). Genes upregulated in *Uhrf1^fl/fl^* ex-FOXP3 cells included some associated with alternate effector T cell function and apoptosis (*Il17rb, Il13, Crtc2, Casp8ap2,* and *Tnfrs8*) **(Supplemental Figure 6D-E and Supplemental Table 1)**. To further elucidate whether engraftment was influenced by the inflammatory microenvironment of the lung, we harvested adoptively transferred *Foxp3*-GFP^+^tdTomato^+^ cells from the spleens of recipients of *Uhrf1^+/+^*and *Uhrf1^fl/fl^* iTregs that received DTx but not influenza and found consistent differences in the frequency of *Uhrf1^+/+^* and *Uhrf1^fl/fl^* iTregs, suggesting a cell-intrinsic defect in engraftment due to loss of UHRF1 **(Supplemental Figure 6F)**. FOXP3 stability was unaffected **(Supplemental Figure 6G)**. Collectively, these data suggest that iTregs require UHRF1 to stabilize their phenotypic identity, upregulate repair processes, and promote tissue engraftment following influenza pneumonia.

**Supplemental Figure 6:**
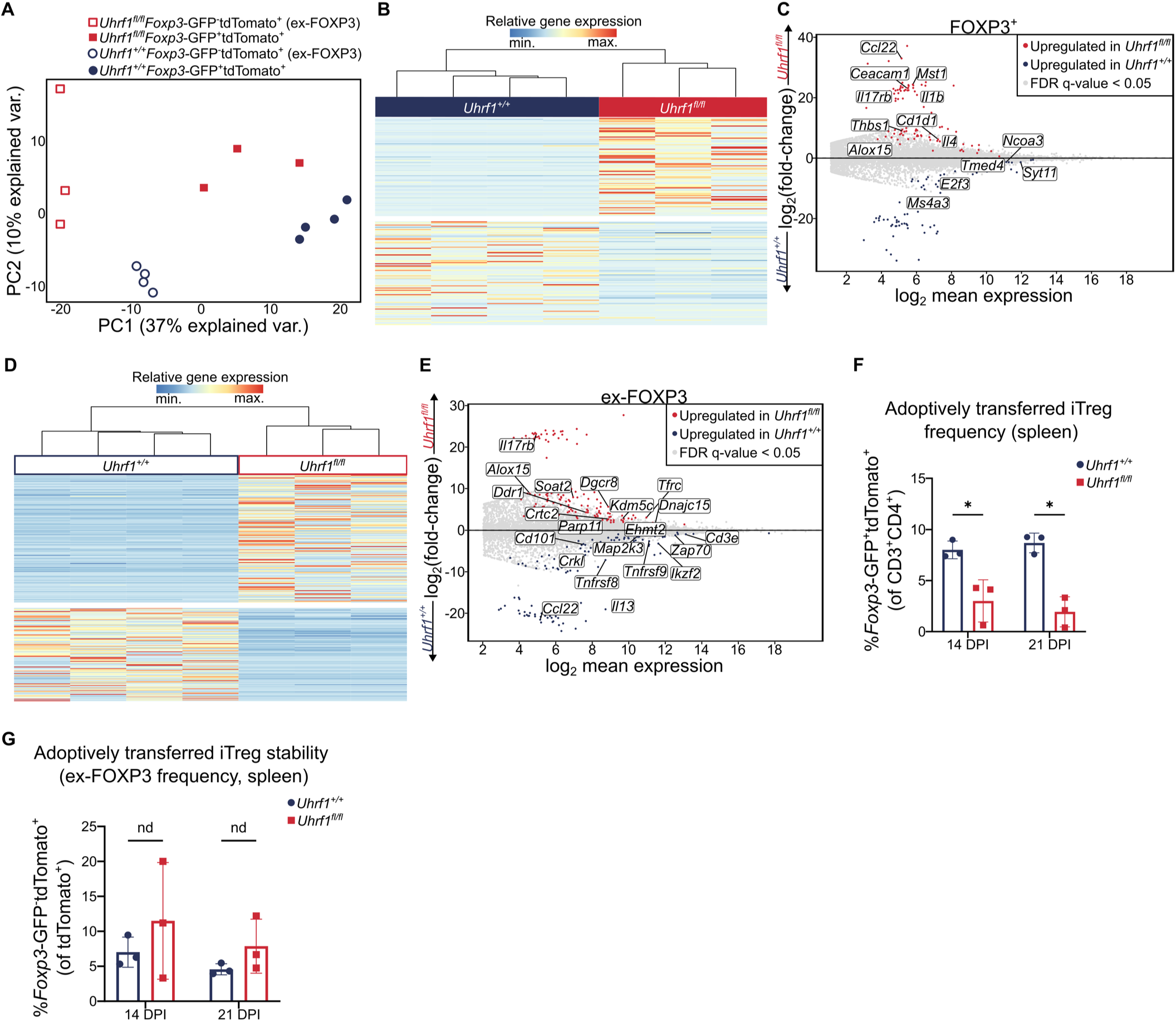
UHRF1 is required for iTreg phenotypic stability and splenic tissue engraftment following viral pneumonia. **(A)** PCA of 457 differential expressed genes from sorted *Uhrf1^+/+^* and *Uhrf1^fl/fl^* cells harvested from the spleen 24 DPI identified from ANOVA-like testing with FDR *q* < 0.05. **(B)** K-means clustering of 183 genes with FDR *q* < 0.05 comparing FOXP3^+^ *Uhrf1^+/+^* and *Uhrf1^fl/fl^* iTregs harvested from the spleens of *Foxp3^GFP-DTR^* mice at 24 DPI with k = 2. **(C)** MA plot comparing gene expression of adoptively transferred FOXP3^+^ *Uhrf1^+/+^* and *Uhrf1^fl/fl^* iTregs harvested from the spleen 24 DPI. Genes of interest are annotated. **(D)** K-means clustering of 274 genes with FDR *q* < 0.05 comparing ex-FOXP3 *Uhrf1^+/+^* and *Uhrf1^fl/fl^* cells harvested from the spleens of *Foxp3^GFP-DTR^* mice at 24 DPI with k = 2. **(E)** MA plot comparing gene expression of adoptively transferred ex-FOXP3 *Uhrf1^+/+^* and *Uhrf1^fl/fl^* cells harvested from the spleen 24 DPI. Genes of interest are annotated. **(F)** Total tdTomato^+^ cells recovered at 14 and 21 DPI from the spleens of recipients of *Uhrf1^fl/fl^* or *iUhrf1^+/+^* iTregs that received DTx but no influenza. **(G)** Frequency of *Foxp3*-GFP^−^tdTomato^+^ cells recovered at 14 and 21 DPI from the spleens of recipients of *Uhrf1^fl/fl^* or *Uhrf1^+/+^* iTregs that received DTx but no influenza. Data presented as mean and SD. * q < 0.05 according to two-way ANOVA with two-stage linear step-up procedure of Benjamini, Krieger, and Yekutieli with Q = 5%. nd, no discovery.

### Loss of UHRF1-mediated maintenance DNA methylation results in disrupted DNA methylation and delayed expression of signature Treg transcriptional programs

To determine transcriptional differences in iTregs earlier in the course of injury and assess for differences in the transcriptional landscape over time, we performed transcriptional profiling of sorted *Uhrf1^fl/fl^* and *Uhrf1^+/+^* cells at 11 DPI. PCA of 2,117 DEGs identified following ANOVA-like testing with FDR *q* < 0.05 demonstrated clustering by DPI and genotype **(Figure 6A)**. PC1 reflected the transcriptional differences between iTregs at 11 versus 24 DPI and PC2 reflected differences between genotypes (*Uhrf1^+/+^* versus *Uhrf1^fl/fl^*) at 24 DPI. Pairwise comparison of cells isolated at 11 DPI revealed 32 DEGs. GSEA revealed positive enrichment of hallmark processes associated with essential Treg functions, including Myc targets, oxidative phosphorylation, E2F targets, TNF-α signaling via NFκB, MTORC signaling, and IL-2–STAT5 signaling in *Uhrf1^+/+^* iTregs. No processes were positively enriched in *Uhrf1^fl/fl^* iTregs **(Figure 6B and Supplemental Table 4)**. Additional GSEA revealed a similar pattern, with positive enrichment of GO processes seen in *Uhrf1^+/+^* iTregs including protein translation, T cell differentiation, regulation of lymphocyte mediated immunity, DNA damage repair, cellular stress responses, and apoptotic processes, but no positively enriched GO processes in *Uhrf1^fl/fl^* iTregs **(Figure 6C and Supplemental Table 4)**. An unsupervised analysis comparing differentially methylated regions (DMRs) of *Uhrf1^fl/fl^* and *Uhrf1^+/+^* iTregs at 24 DPI with at least a 10% difference in methylation demonstrated disrupted methylation at 34 regions in *Uhrf1^fl/fl^* iTregs **(Figure 6D and Supplemental Table 1)**. These findings fit a pattern in which processes enriched in *Uhrf1^+/+^* iTregs at 11 DPI are not enriched in *Uhrf1^fl/fl^* iTregs until 24 DPI, suggesting a delayed transcriptomic phenotype paralleling the delayed repair phenotype of the recipient mice.

**Figure 6:**
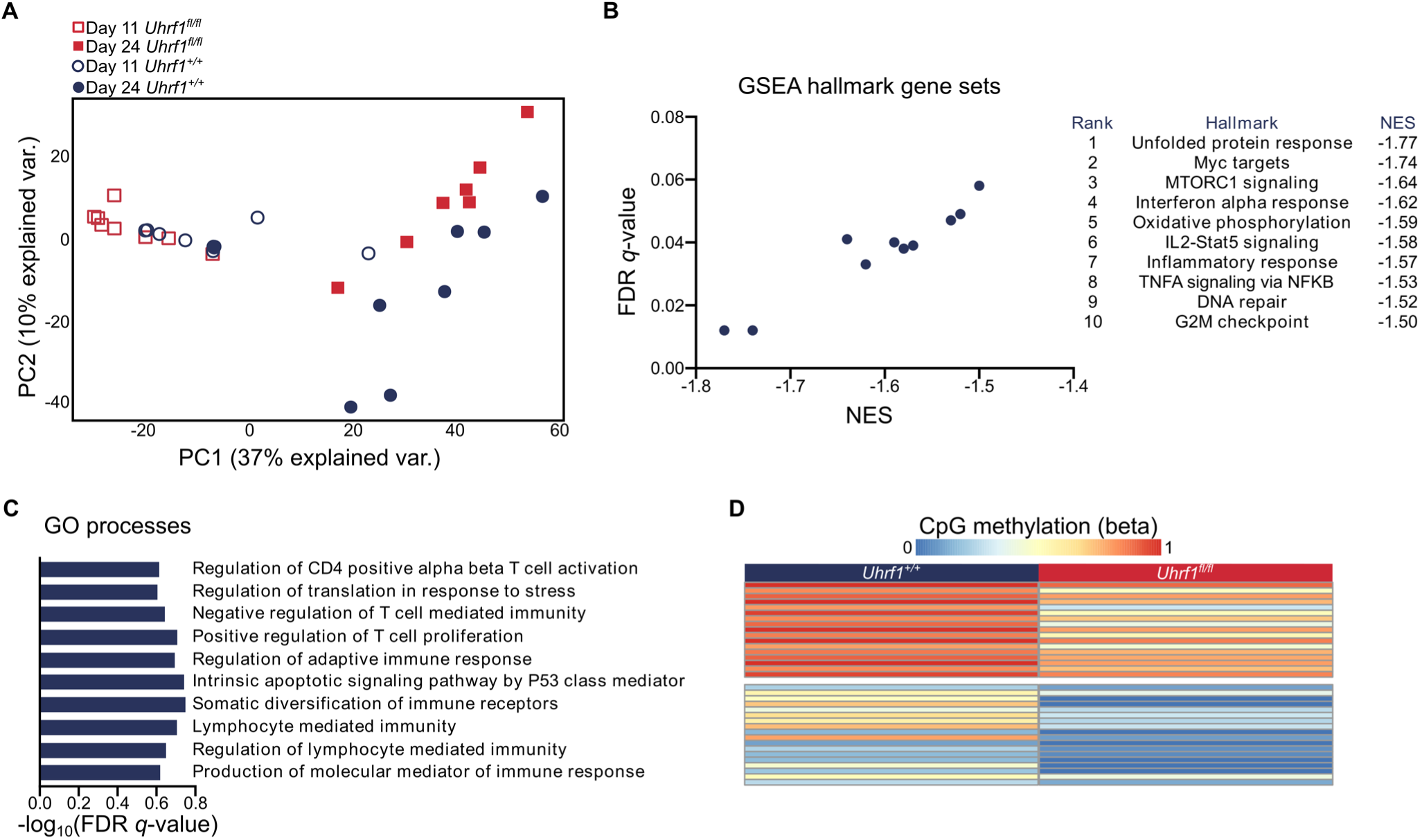
Loss of UHRF1 results in delayed transcriptional changes over the course of viral pneumonia. **(A)** PCA of 2,117 differential expressed genes in sorted *Uhrf1^+/+^*and *Uhrf1^fl/fl^* iTregs from the lungs of recipient *Foxp3^GFP-DTR^*mice at 11 and 24 DPI, identified from ANOVA-like testing with FDR *q* < 0.05. **(B)** GSEA dot plot highlighting key statistics (FDR *q*-value and normalized enrichment score or NES) and enriched gene sets for *Uhrf1^+/+^* iTregs at 11 DPI. Blue dots denote gene sets with a negative enrichment score or enrichment at the bottom of the ranked list. **(C)** Selection of representative enriched gene ontology (GO) processes from 40 total enriched gene sets with an FDR *q* < 0.25 in *Uhrf1^+/+^* iTregs at 11 DPI. Gene sets are annotated and ranked by – log_10_-transformed FDR *q* value. **(D)** K-means clustering of 34 DMRs with a difference of ≥10% between *Uhrf1^+/+^* and *Uhrf1^fl/fl^* iTregs.

## Discussion

Natural Tregs depend on specific patterns of DNA methylation to establish cell identity and stability as well as to exert their pro-recovery function following acute lung injury (8,12,30,48). Here, we demonstrated the ability of transferred iTregs to promote recovery following viral pneumonia in mice. We further demonstrated that maintenance DNA methylation mediated by the epigenetic regulator, UHRF1, is necessary for iTreg reparative function during viral pneumonia. Recipients of UHRF1-deficient iTregs experienced worsened hypoxemia and mortality with dysregulated and delayed lung repair. In vitro, the loss of maintenance DNA methylation resulted in downregulation of chemokine receptors, such as CCR4, CCR5, and CCR8. In vivo, UHRF1-deficient iTregs displayed reduced lung tissue engraftment as well as downregulation of genes associated with pro-repair function and upregulation of pro-apoptotic genes and alternate effector T cell lineage defining transcription factors. These findings support the use of iTregs as for cellular therapy while demonstrating the necessity of maintenance DNA methylation in iTreg stability and function.

Our observations support a paradigm in which critical changes occur across the DNA methylation landscape between the CD4^+^ stage of T cell development and subsequent FOXP3 expression that cause a differential effect of the loss of maintenance DNA methylation on iTreg function. We demonstrated that iTregs that lose UHRF1 following FOXP3 induction possess a reduced ability to protect the lung from alveolar epithelial injury, leading to delayed lung repair. In contrast, iTregs generated from UHRF1-deficient CD4^+^ T cells demonstrated hyper-suppressive function when adoptively transferred into lymphocyte-deficient mice with Tconv cell-mediated colitis (31). In both cases, the loss of UHRF1 did not affect FOXP3 induction or stability but did alter the transcriptome at other loci. A similar effect dependent on the timing of loss of UHRF1 exists in nTregs. Work from our group demonstrated that the loss of UHRF1 at the FOXP3+ stage of nTreg development caused the spontaneous onset of widespread scurfy-like inflammation (30). Intriguingly, in that study, we also observed the generation of ex-FOXP3 cells following the induced loss of UHRF1 in vivo, suggesting a link between UHRF1-mediated maintenance DNA methylation and the stability of FOXP3 expression in nTregs. In contrast, mice with pan-T cell UHRF1 deficiency develop inflammation only localized to the colon (49). Although the latter study noted differences in thymic and peripheral Treg populations, it further serves as evidence of the differential effect of the loss of UHRF1 depending on the developmental stage of Tregs.

Despite reports of instability of iTregs within inflammatory microenvironments, our data suggest that iTregs retain their function in promoting lung repair similar to adoptively transferred nTregs (12,16–18). Suppressive mechanisms did not appear to play a role in the differential effects of the loss of UHRF1, as immune infiltration and activation were similar in mice that received UHRF1-sufficient and -deficient iTregs. Whether and how the inflammatory microenvironment contributed to reduced engraftment of UHRF1-deficient iTregs is unclear. RNA sequencing data from cells cultured in vitro suggest that reduced engraftment may have been a result of reduced homing receptors, such as CCR4, 5, and 8. In addition, data from UHRF1-deficient iTregs recovered from the lung revealed upregulation of several pro-apoptotic factors, suggesting an influence from the inflammatory microenvironment that reduced UHRF1-deficient iTreg numbers.

iTregs may therefore serve as a practical, safe, and efficient alternative to nTreg cellular therapy for severe, rapidly progressive inflammatory diseases such as ARDS (50). Currently, nTregs are obtained via leukapheresis from autologous peripheral or allogeneic cord blood, and protocols to expand them are on the order of weeks (51,52). This same process could be applied to iTregs in a fraction of the time, as they derive from CD4^+^ Tconv cells and expand rapidly in culture, allowing for intervention at an earlier time point in the ARDS disease course. Similar to strategies for nTregs, ex vivo modification strategies could be implemented to further enhance iTreg therapeutic efficacy (15). As we noted that cellular engraftment may play a role in the differential effects we observed in recipients of UHRF1-sufficient and -deficient iTregs, in vitro supplementation with factors to promote homing to the site of injury via enhanced chemokine receptor expression could ensure cellular presence at the site of injury. Additionally, as the loss of UHRF1-mediated maintenance DNA methylation resulted in upregulation of genes associated with effector helper T cell lineage commitment, factors to promote their downregulation could be leveraged to ensure stabile functioning.

In summary, our data establish that iTregs promote timely repair of damaged lung tissue following viral pneumonia. Mechanistically, UHRF1-mediated maintenance DNA methylation is required for optimal iTreg engraftment and reparative function. These data credential iTregs as potential cellular therapy to promote repair following viral pneumonia-induced ARDS.

## Methods

### Mice

*Foxp3^GFP-DTR^* mice were purchased from The Jackson Laboratory (strain no. 016958) and bred on-site. C57BL/6 *Uhrf1^fl/fl^Foxp3^GFP-CreERT2^Rosa26Sor^CAG-tdTomato^*or *Foxp3^GFP-CreERT2^Rosa26Sor^CAG-tdTomato^* mice were generated as previously described (30). All animals were genotyped using services provided by Transnetyx Inc., with primer sequences provided by The Jackson Laboratory or published in prior work (30). Animals received water ad libitum, were housed at a temperature range of 20 °C to 23 °C under 14-hour light/10-hour dark cycles and received standard rodent chow.

### Tissue preparation, flow cytometry, and cell sorting

Spleens and lymph nodes (inguinal and axillary) were harvested from adult (8–12-week-old) mice. To obtain a single-cell suspension, tissues were disrupted using scored 60-mm petri dishes in PBS and filtered through a 40-μm nylon mesh filter. Red blood cell lysis was performed using Gibco ACK lysing buffer (cat. no. A1049201). Cell counts were obtained using a Cellometer K2 Counter using AOPI stain (Nexcelom Bioscience cat. no. SD014-0106). CD4^+^ T cells were purified from single-cell suspension using the EasySep™ mouse CD4^+^ T cell Isolation Kit (Stemcell, cat. no. 19852) or the negative fraction of the Miltenyi mouse CD4^+^CD25^+^ Regulatory T cell Isolation Kit (cat. no. 130-091-041) according to the manufacturer’s instructions. Enriched CD4^+^ single-cell suspensions were stained with antibodies against CD4 prior to flow cytometry sorting (**Supplemental Table 5**). To further enhance purity prior to culture for iTreg induction, conventional CD4^+^ T cells were separated from CD4^+^*Foxp3*-GFP^+^ nTregs with a microfluidics MACSQuant® Tyto® sorter (Miltenyi). Dead cells were excluded using a viability dye for analysis and sorting (**Supplemental Table 5**) in all experiments. For Treg adoptive transfer, natural and induced Tregs were harvested from splenic and lymph node single-cell suspension or culture, respectively, re-stained for CD4 and viability dye, and sorted using the MACSQuant® Tyto® sorter (Miltenyi) (**Supplemental Table 5**).

**Supplemental Table 5.** Flow cytometry fluorochromes and reagents.

| Antigen/Reagent | Conjugate | Clone | Manufacturer | Catalog no. |
| --- | --- | --- | --- | --- |
| CD4 | APC-eFluor™ 780 | RM4-5 | Invitrogen | 47-0042-82 |
| DAPI/Cell-Impermeant Dye | N/A | N/A | Thermo Fisher Scientific | 62248 |
| CD45 | FITC | 30-F11 | BioLegend | 103107 |
| CD31 | PE | MEC 13.3 | BD Pharmingen | 553373 |
| Podoplanin | PE-Cyanine7 | eBio8.1.1 | eBioscience | 25-5381-82 |
| CD326 | BV421 | G8.8 | BioLegend | 118225 |
| Krt5; Krt5 Secondary Antibody | Unconjugated Alexa Fluor 488 | Poly9059 | BioLegend; Invitrogen | 905901; A11039 |
| MHCII | BUV395 | 2G9 | BD OptiBuild | 743876 |
| Ki-67 | PerCP-eFluor 710 | SolA15 | eBioscience | 46-5698-82 |
| Fixable Viability Dye eFluor 506 | N/A | N/A | eBioscience | 65-0866-14 |
| CD45 | APC-Cy7 | 30-F11 | BD Pharmingen | 557659 |
| CD3e | FITC | 145-2C11 | Invitrogen | 11-0031-85 |
| CD4 | BUV395 | GK1.5 | BD Horizon | 565974 |
| CD8 | PECF594 | 53-6.7 | BioLegend | 100762 |
| CD64 | PE | X54-5/7.1 | BioLegend | 139304 |
| CD11b | BUV737 | M1/70 | BD Horizon | 612800 |
| CD11c | Pe-Cy7 | HL3 | BD Pharmingen | 558079 |
| SiglecF | AF647 (APC) | E50-2440 | BD Pharmingen | 562680 |
| Ly6G | AF 700 (A700) | 1A8 | BD Pharmingen | 561236 |
| CountBright™ Absolute Counting Beads | N/A | N/A | Invitrogen | C36950 |

### Cell culture and iTreg induction

For nTreg culture, 100,000 CD4^+^*Foxp3*-GFP^+^ Tregs were seeded in 96-well round bottom plates (Corning ref. no. 3799) with αCD3/αCD28 Dynabeads™ (Gibco ref. no. 11456D) at a ratio of three beads to one Treg cell and recombinant human IL-2 at a concentration of 2,000 U/ml (NCI Frederick National Laboratory). For iTreg culture, 300,000 CD4^+^*Foxp3*-GFP^−^ conventional T cells were seeded in 24-well plates (Fisher ref FB012929) coated with αCD3/αCD28 antibody (BD Pharmingen cat. no. 553058 and 553295) at concentrations of 3 µg/ml and 5 µg/ml, respectively. Recombinant human IL-2 and TGF-β (PeproTech cat. no. 100-21-10UG) were added at 50 U/ml and 10 ng/ml, respectively. Both cell types were cultured in RPMI 1640 (Gibco cat. no. 11875-093) media supplemented with 10% fetal bovine serum, 5 mM HEPES (Gibco cat. no. 15630080), 100 U/ml penicillin-streptomycin (Gibco cat. no. 15140122), 1 mM Na-pyruvate (Gibco cat. no. 11360-070), 100 mM MEM non-essential amino acids (Gibco cat. no. 11140050), 2 mM L-glutamine (Gibco cat no. 25030081), and 50 µM 2-mercaptoethanol (Sigma-Aldrich cat. no. M3148). For FOXP3^+^ cell lineage tracing and *Foxp3*-Cre-mediated loss of UHRF1, Z-4-hydroxytamoxifen (Sigma-Aldrich cat. no. H7904) was added to cell culture at a concentration of 500 nM.

### Suppression assays

For suppression assays, CD4^+^*Foxp3*-GFP^+^ Tregs were co-cultured with varying ratios of CD4^+^*Foxp3*-GFP^−^ conventional T cells labeled with Cell Trace Violet (Invitrogen ref. no. C34557) according to the manufacturer’s instructions and as previously described (40). Treg-CD4^+^ conventional T cell co-cultures were activated using αCD3/αCD28 Dynabead™ particles at a ratio of three beads to one Treg cell. After 72 hours, cells were harvested and analyzed by flow cytometry using a BD LSRFortessa. The division index of each sample was calculated using the proliferation modeling function in FlowJo software version 10.

### Diphtheria toxin and influenza A virus administration

To ablate the endogenous mouse Treg population, lyophilized diphtheria toxin (List Biological Laboratories, product no. 150) was resuspended in sterile PBS and administered in 100 µl via intraperitoneal injection every two days to *Foxp3^GFP-DTR^* mice. The initial loading dose was 50 µg/kg followed by maintenance dosing of 10 µg/kg. For influenza A virus administration, mice were intubated while under isoflurane anesthesia using a 20-gauge angiocatheter cut to a length that placed the tip of the catheter above the carina. The mice were then instilled with mouse-adapted influenza A/WSN/33 H1N1 virus in 50 μL of sterile PBS as previously described (33).

### Measurement of physiologic readouts of viral pneumonia progression and recovery

To prepare mice for arterial oxyhemoglobin saturation (SpO_2_) measurement, one day prior to influenza inoculation, a depilatory cream containing the active ingredients potassium thioglycolate (4%) and calcium hydroxide (1.5%) (Nair^TM^) was applied to the hair on the dorsal neck for two minutes and subsequently gently removed with wet gauze under isoflurane anesthesia. A MouseOx Plus pulse oximeter (Starr Life Sciences) was used to measure SpO_2_. Unanesthetized mice were immobilized, and an oximeter collar clip was secured to the hairless neck. Baseline SpO_2_ and mass measurements were obtained prior to influenza administration with subsequent measurements taken every other day post-inoculation.

### Adoptive transfer

Prior to adoptive transfer, cells were harvested from culture and sorted via a microfluidics MACSQuant® Tyto® sorter (Miltenyi) to enhance purity. Cells were subsequently washed twice and resuspended in sterile PBS at a concentration of 1×10^6^ cells/100 µl. Once resuspended, cells were drawn up into a 500-µl insulin syringe (BD ref 329461) and delivered into mice placed under isoflurane anesthesia via the retroorbital sinus (40).

### Lung tissue harvesting, processing, and analysis

Mice were euthanized using a carbon dioxide euthanasia chamber followed by cervical dislocation and slowly infused with HBSS through the right atrium of the heart to clear the pulmonary circulation of blood. The lungs were then harvested and the larger airways and other mediastinal structures were trimmed. The post-caval lobes were removed and set aside for epithelial cell analysis by flow cytometry. For adoptively transferred iTreg and infiltrating immune cell analysis, the remaining lung lobes were grossly homogenized with scissors in HBSS containing 2 mg of collagenase D (Sigma Aldrich cat. no. 11088866001) and 0.25 mg of DNAse I (Sigma Aldrich cat. no. 10104159001) per ml. The lung suspensions were subsequently incubated for 45 minutes at 37 °C and then further homogenized using a Miltenyi OctoMACS tissue dissociator using the mouse lung protocol (m_lung_02). For iTreg cell sorting, the single-cell suspension was enriched using a CD4^+^ antibody (Miltenyi cat. no. 130-101-962) and the lungs were stained using the reagents listed in **Supplemental Table 5**. A separate aliquot was taken for infiltrating immune cell analysis, which was enriched using CD45^+^ selection beads (Miltenyi cat. no. 130-052-301) and the cells were stained using the reagents listed in **Supplemental Table 5**. Mice were included in the final analyses if their lungs displayed evidence of gross injury or they developed desaturation to at least 89%, indicating successful influenza infection.

For post-influenza induced lung injury epithelial repair analysis, the harvested post-caval lobes were injected with 2 mL Dispase (Corning ref. 354325) containing 0.25 mg DNAse I per mL and incubated for 45 minutes at room temperature on a rocker. Afterward, forceps were used to tease apart large pieces of lung tissue followed by more thorough homogenization via vigorous pipetting through wide-bore pipette tips. The homogenized tissue was then incubated for another 10 minutes at room temperature on a rocker and then filtered through 40-µm filters. Following centrifugation, the cells were resuspended in 2 mL ACK RBC lysis buffer and incubated for an additional four minutes at room temperature. The remaining cells were fixed and stained for flow cytometry analysis using the reagents listed in **Supplemental Table 5**. Data acquisition for analysis was performed using a Symphony A5 instrument with FACSDiva software (BD). Analysis was performed with FlowJo software version 10.

For preparation of lungs for histopathology, mice were euthanized via carbon dioxide euthanasia chamber and a tracheostomy was created. A 20-gauge angiocatheter was subsequently sutured into the trachea via the tracheostomy. The lungs and trachea were removed en bloc and the lungs were inflated to 15 cm H_2_O with 4% paraformaldehyde. 5-μm sections from paraffin-embedded lungs were stained with H&E or trichrome and examined using light microscopy with a high-throughput, automated slide imaging system, TissueGnostics (TissueGnostics GmbH).

### RNA-sequencing and analysis

Adoptively transferred iTregs isolated from the spleen post influenza or from in vitro cultured cells were harvested from splenic single-cell suspension or cell culture, respectively, flow cytometry sorted, and lysed immediately after sorting with QIAGEN RLT Plus (cat. no. 1053393) containing 1% 2-mercaptoethanol. Cells were then subjected to simultaneous RNA and DNA isolation using the QIAGEN AllPrep Micro Kit (cat. no. 80204). RNA-library preparation was performed using the SMARTer Stranded Total RNA-Seq Kit, version 2 (Takara cat. no. 634411) as previously described (30,53,54). Sequencing was performed on an Illumina NextSeq 2000 instrument as previously described (30). For rare adoptively transferred iTregs, cells were flow cytometry sorted following harvest from lung single-cell suspensions and lysed immediately after sorting with 10x RNA lysis buffer containing 5% RNase inhibitor. 50% of the sample was taken for DNA isolation using the QIAGEN AllPrep Micro Kit (cat. no. 80204). RNA-seq libraries were then prepared by using 100 pg total RNA from each sample by following the SMART-Seq v4 Ultra Low Input RNA Kit (Takara Bio user manual. The cDNA was amplified with 11 cycles of PCR. The Nextera XT kit (Nextera XT DNA Library Preparation Kit, Illumina) was used to make cDNA libraries suitable for Illumina sequencing. Prepared libraries were pooled at 4 nM and sequenced on a NextSeq 2000 (Illumina) using 75-base read lengths in single-end mode.

RNA-seq analysis was performed as previously described (55). After sequencing, raw binary base call (BCL) files were converted to FASTQ files using bcl-Convert (version 3.10.5, Illumina). Adaptor trimming, alignment to the GRCm38 reference genome, and quantification were performed using the nf-core/RNA-seq pipeline version 3.9 (implemented in Nextflow 22.04.5 with Northwestern University Genomics Compute Cluster configuration (nextflow run nf-core/rnaseq -profile nu_genomics --genome GRCm38). Differential expression analysis was performed in R package DEseq2 (version 1.38.3 in R 4.2.3). For K means analysis, k was determined using elbow plots and the kmeans function in R stats 3.6.2 (Hartigan–Wong method with 25 random sets and a maximum of 1,000 iterations) was used for clustering. In vitro K means heat maps generated using the Morpheus web interface (https://software.broadinstitute.org/morpheus/). Gene Set Enrichment Analysis was performed using the Broad Institute’s GSEA software, version 4.1.0, GSEAPreranked tool (56) with genes ordered by log_2_(fold-change) in average expression against the Hallmark gene sets or the Immunologic Signature gene sets housed in the Molecular Signatures Database of the Broad Institute (57).

### Modified reduced representation bisulfite sequencing (mRRBS) and reduced representation enzymatic methylation sequencing (RREM-seq) and analysis

mRRBS library preparation for in vitro cultured cells was performed using procedures previously described by our group (30,40,58). Briefly, genomic DNA was isolated from sorted samples via the QIAGEN AllPrep Micro Kit (cat. no. 80204) and quantified with a Qubit 3.0 instrument. Bisulfite conversion was then performed using the EZ DNA Methylation-Lightning Kit (Zymo Research) per the manufacturer’s protocol. We next created indexed Illumina-compatible non-directional libraries from bisulfite-converted single-stranded DNA using the Pico Methyl-Seq Library Prep Kit (Zymo Research). Final library size distribution and quality were assessed via high-sensitivity screen tape (TapeStation 4200, Agilent). Libraries were then pooled for sequencing on a NextSeq 2000 (Illumina) instrument using the V2 High Output reagent kit (1 × 75 cycles).

RREM-seq library preparation was performed as previously described (59,60). Briefly, following genomic DNA extraction using the QIAGEN AllPrep DNA/RNA Micro Kit (cat. no. 80204) 0.5–1 ng of genomic DNA was digested using restriction endonuclease MspI (New England Biolabs) and then enzymatically converted with TET2 and APOBEC (New England Biolabs) per the manufacturer’s instructions. Random priming, adapter ligation, PCR product clean-up, and final library amplification were performed using the Pico methyl-seq library prep kit (Zymo Research). Unmethylated λ-bacteriophage DNA (1:200 mass ratio; New England Biolabs) was included in all samples to calculate unmethylated cytosine conversion efficiency (on average >99%). Final library size distribution and quality were assessed via high-sensitivity screen tape (TapeStation 4200, Agilent) and sequenced using single-end reads with a NextSeq 2000 (Illumina).

Methylation analysis was conducted as previously described (30,59). Briefly, following sequencing raw binary base call (BCL) files were converted to FASTQ files using bclConvert (version 3.10.5, Illumina) and trimmed using Trim Galore! (version 0.4.3). Bismark (version 0.21.0) was used to perform alignment to the reference genome mm10 (GRCm38) and methylation extraction. Bismark coverage files were used for quantification using SeqMonk (version 1.48.0) and R package DSS (version 2.46.0). Cumulative distribution plots were generated with the ecdf base R function.

### Statistics

*P*-values and *q*-values resulting from two-tailed tests were calculated using statistical tests stated in the figure legends. Statistical analysis was performed using either GraphPad Prism version 10.3.0 or R version 4.2.3. A *p*- or *q*-value of less than 0.05 was considered significant except for GSEA, in which 0.25 was considered significant (56). Outliers were identified and removed via the ROUT method at a Q of 5%. Computational analysis was performed using Genomics Nodes and Analytics Nodes on Quest, Northwestern University’s High-Performance Computing Cluster.

### Study approval

All animal experiments and procedures were conducted in accordance with the standards established by the US Animal Welfare Act set forth in NIH guidelines and were approved by the IACUC at Northwestern University under protocols IS00012519 and IS00017837.

### Data and material availability

Raw sequencing data will be made publicly available on the GEO database pending peer-reviewed publication.

## Author contributions

AMJ, JKG, MATA, SEW, and BDS contributed to the conception, hypothesis delineation, and design of the study. AMJ, JKG, QL, MATA, KAH, LM-N, NM, CPRF, HAV, EMS, SEW, and BDS performed experiments/data acquisition and analysis. AMJ and BDS wrote the manuscript or provided substantial involvement in its revision.

## Acknowledgements

AMJ is supported by NIH award F32HL162418. MATA is supported by NIH awards T32GM144295, T32HL076139, and F31HL162490. NM is supported by NIH award T32AI083216. CPRF is supported by T32HL076139. JG is supported by T32HL076139. QL is supported by the David W. Cugell Fellowship and the The Genomics Network (GeNe) Pilot Project Funding. LM-N is supported by the NIH awards K08HL159356, U19AI135964 and the Parker B. Francis Opportunity Award. SEW is supported by the Burroughs Wellcome Fund Career Awards for Medical Scientists. BDS is supported by NIH awards R01HL149883, R01HL153122, P01HL154998, P01AG049665, U19AI135964, and U19AI181102. We wish to acknowledge the Northwestern University Flow Cytometry Core Facility supported by CA060553; the BD FACSAria SORP system was purchased with the support of S10OD011996. We also wish to acknowledge the Northwestern University RNA-Seq Center/Genomics Lab of the Pulmonary and Critical Care Medicine and Rheumatology Divisions. Histology services were provided by the Northwestern University Mouse Histology and Phenotyping Laboratory, which is supported by P30CA060553 awarded to the Robert H. Lurie Comprehensive Cancer Center. This research was supported in part through the computational resources and staff contributions provided by the Genomics Compute Cluster, which is jointly supported by the Feinberg School of Medicine, the Center for Genetic Medicine, and Feinberg’s Department of Biochemistry and Molecular Genetics, the Office of the Provost, the Office for Research, and Northwestern Information Technology. The Genomics Compute Cluster is part of Quest, Northwestern University’s high performance computing facility, with the purpose to advance research in genomics. The graphical abstract, Figure 1A, Supplemental Figure 1A, and Figure 2A were created using https://www.biorender.com/.

